# Comparison of Three Transcytotic Pathways for Distribution to Brain Metastases of Breast Cancer

**DOI:** 10.1101/2022.08.09.503253

**Authors:** Imran Khan, Brunilde Gril, Anurag Paranjape, Christina Robinson, Simone Difilippantonio, Wojciech Biernat, Michał Bieńkowski, Rafał Pęksa, Renata Duchnowska, Jacek Jassem, Priscilla K. Brastianos, Philippe Metellus, Emilie Bialecki, Carolyn C. Woodroofe, Haitao Wu, Rolf Swenson, Patricia S. Steeg

## Abstract

Advances in drug treatments for brain metastases of breast cancer have improved progression free survival but new, more efficacious strategies are needed. Most chemotherapeutic drugs infiltrate brain metastases by moving between brain capillary endothelial cells, paracellular distribution, resulting in heterogeneous distribution, lower than that to systemic metastases. Herein, we tested three well-known transcytotic pathways through brain capillary endothelial cells as potential avenues for drug access: Transferrin receptor (TfR) peptide, Low density lipoprotein receptor 1 (LRP1) peptide, Albumin. Each was far-red labeled, injected into two hematogenous models of brain metastases, circulated for two different times, and their uptake quantified in metastases and uninvolved (nonmetastatic) brain. Surprisingly, all three pathways demonstrated distinct distribution patterns *in vivo.* Two were suboptimal: TfR distributed to uninvolved brain but poorly in metastases, while LRP1 was poorly distributed. Albumin distributed to virtually all metastases in both model systems, significantly greater than in uninvolved brain (P <0.0001). Further experiments revealed that albumin entered both macrometastases and micrometastases, the targets of treatment and prevention translational strategies. Albumin uptake into brain metastases was not correlated with the uptake of a paracellular probe (biocytin). We identified a novel mechanism of albumin endocytosis through the endothelia of brain metastases consistent with clathrin-independent endocytosis (CIE), involving the neonatal Fc receptor (FcRn), galectin-3 (Gal-3) and glycosphingolipids. Components of the CIE process were found on metastatic endothelial cells in human craniotomies. The data suggest a reconsideration of albumin as a translational mechanism for improved drug delivery to brain metastases and possibly other CNS cancers.

**Statement of Significance:** Drug therapy for brain metastasis needs improvements. We surveyed transcytotic pathways as potential delivery systems in brain-tropic models and found that albumin has optimal properties. Albumin used a novel mechanism for endocytosis.

## Introduction

Brain (CNS) metastases of breast and lung cancers and melanoma confer serious physical and neurocognitive effects and contribute to patients’ deaths (rev. in (1–3)). Patients with two subtypes of metastatic breast cancer, tumors with HER2 amplification/overexpression (HER2+), and tumors that are hormone receptor negative, HER2 normal (triple-negative) are at highest risk. Standard treatments include surgery, stereotactic radiation therapy to the lesions, whole brain radiation therapy, and steroids for edema. Drug therapies for breast cancer brain metastases have a checkered history. The most brain permeable drugs, such as molecular inhibitors of HER2, offer only months of increased progression free- or overall survival (4,5). Many other drugs, however, even those with known systemic activity in the metastatic setting were virtually inactive in the CNS (6,7). In agreement, drug distribution to brain metastases in hematogenous model systems is heterogeneous (8–13) and remains ~a log below that of systemic metastases (11).

The limited efficacy of drugs for CNS metastases is a major hurdle to advances in patient care. In addition to normal pharmacokinetic parameters of metabolism, protein binding, excretion, edema etc., drug distribution to brain metastases necessarily involves traversal through the blood-tumor barrier (BTB), an alteration of the normal capillary blood-brain barrier (BBB) that protects brain parenchyma (rev. in (14)). Most drugs studied to date traverse the BTB paracellularly: Endothelial cells of the normal BBB are connected by continuous tight- and adherens junctions which break down to some extent in the BTB allowing drug passage between cells.

An alternative route of drug distribution to brain metastases is transcytosis, through the capillary endothelia and other cells. Transcytosis is a complex series of mechanisms for the intracellular transport of macromolecules within membrane-bound vesicles. In the normal brain needed metabolites enter via receptor-mediated transport, carrier-mediated transport, lipid transport, lipid diffusion and, to a very limited extent, macropinocytosis of soluble compounds (rev. in (15,16)). Transcytotic pathways vary in initial modes of cell membrane endocytosis (clathrin-coated pits, caveolae, clathrin-independent endocytosis, etc.). Intracellular vesicles can route to lysosomal degradation, recycle back to the cell membrane, or move across the cell to release their contents.

Several drugs connected to transcytotic pathway ligands have been preclinically tested in either the normal brain, non-cancer CNS diseases such as Alzheimers’, primary CNS malignancies or brain metastases. We have little research to explain disparate results, which would guide future development efforts. For instance, the transferrin receptor (TfR) pathway delivers iron to the normal brain via receptor mediated transcytosis and has been linked to a number of drugs or used to coat nanoparticles (rev. in (17)). Antibodies to TfR have shown brain transcytosis that is dependent on antibody avidity in Alzheimers’ studies (18); in normal brain, conjugation of TfR antibodies or Tf peptides bound to quantum dots yielded low brain distribution, with the former trapped in endothelial cells (19,20). To our knowledge, TfR-based investigations in brain metastases have not been reported. Another transcytotic pathway investigated is the low density lipoprotein receptor related protein 1 (LRP1) which initiates endothelial transcytosis and is critical for Tau uptake and spread in neurodegenerative diseases (21). A clinical trial of Angiopep, a LRP1 peptide conjugated to paclitaxel, showed some disease benefit but the trial missed its primary endpoint (22). In a brain metastasis model system Angiopep conjugated to paclitaxel resulted in a 161-fold increase in drug levels compared to free drug in uninvolved brain, but only <10-fold increase in experimental hematogenous metastases (12), suggesting a propensity for normal brain transcytosis over brain metastasis. Another investigation used LRP peptide-conjugated liposomes in an intracranial xenograft where membrane fluidity was a significant factor in controlling intracranial tumor growth (23). A third pathway, albumin, has also been used to coat nanoparticles, either temporarily to improve solubility or in a more stable manner (rev in (24)); some preclinical efficacy in primary brain tumor and brain metastasis models has been reported (25,26). The transcytotic pathway for albumin in normal or diseased brain is unresolved and could include binding to the endothelial neonatal Fc receptor (FcRn)(27) or other proteins including SPARC(28), or entry via fluid filled large pinocytotic vesicles (29,30).

These studies differ by transcytotic pathway, associated drugs, and formulations. In the current manuscript we have attempted to identify an optimal transcytotic pathway in brain metastasis models, which can then be further developed to optimize drugs and formulations. Questions resolved herein include: (a) are all transcytotic pathways equally capable of delivering a ligand to a brain metastasis? (b) Conversely, to what extent does each pathway traverse metastases versus uninvolved brain? Uptake of a toxic drug by uninvolved brain could result in adverse effects. (c) Do transcytotic pathways enable ligand distribution into both macrometastases and micrometastases? The ability to penetrate micrometastases may be fundamental to brain metastatic prevention efforts. Our data in two model systems at two timepoints show an unexpected clear superiority of albumin. With this finding a second major gap in our research knowledge appeared: If albumin preferentially transcytoses through the BTB via fluid filled pinocytotic vesicles, why don’t other drugs pass via the same mechanism? We have resolved this question, developing evidence that albumin uses a protein-specific macropinocytotic endocytic pathway with features of clathrin-independent endocytosis (CIE), involving the FcRn, galectins and membrane lipids. Finally, we demonstrate the relevance of this transcytotic pathway model data demonstrating CIE proteins in human breast cancer brain metastases craniotomy specimens. The data provide a fundamental mechanistic background for translational efforts to improve drug penetration and efficacy for brain metastases and possibly other CNS malignancies.

## Materials and Methods

### Cell line origins and authentication

The origins and validation of the triple-negative 231-BR (31) and the HER2-overexpressing JIMT1-BR (32) are previously reported (see Supplementary methods). For *in vitro* BBB and BTB assays, immortalized human astrocytes (HAL) and pericytes (HPL) were generated using SV40 Large T antigen (33). The mouse brain endothelial line bEnd.3 was used. Culture conditions are listed in Supplementary methods.

### Animal Experiments-General Information

All animal experiments were performed under the regulation of the Animal Care and Use Committee (ACUC) of the National Cancer Institute (NCI). Five-seven week old female athymic NIH nu/nu mice were injected in the left cardiac ventricle with 1.75 x 10^5^ 231-BR or JIMT1-BR cells as described, with metastasis formation in 3-4 weeks (32). Upon sectioning, brain metastases were divided into macrometastases, longer than 300 μm in a single dimension, comparable in size to a 3 mm MRI-detectable lesion in a human; micrometastases were smaller lesions. Further details are in Supplementary methods.

### Comparison of three transcytosis ligands

Three known transcytosis ligands, similarly labeled, were compared for distribution to metastases *in vivo.* After the development of brain metastases and on the day of necropsy, mice were injected IV with a far-red fluorescent probe of either albumin, transferrin peptide or LRP-1 peptide; the probes circulated for a short (0.5-1h) or longer (4-6h) period, followed by perfusion. Whole brain sections were scanned for far-red and eGFP fluorescence using a Zeiss AxioScan.Z1 Slide Scanner. Mean fluorescence intensity of each probe in uninvolved (metastasis free, U) brain and brain metastases (M) were quantitated by ZEN 2 software.

### Comparison of transcytosis versus paracellular permeability

231-BR and JIMT1-BR cells were injected into mice and developed brain metastases. After brain metastasis formation and on the day of necropsy far-red-Albumin (Alexa Fluor^™^ 647-albumin) and Biocytin-TMR was injected (IV) and circulated for 60 and10 min, respectively, followed by the perfusion. Whole brain sections were scanned for far-red, red and eGFP fluorescence or H&E staining using a Zeiss AxioScan.Z1 Slide Scanner. Mean fluorescence intensity of each probe was quantitated by ZEN 2 software.

### *In vitro* blood-brain barrier (BBB) and blood-tumor barrier (BTB) assays

The *in vitro* BBB/BTB assays reflecting paracellular permeability was previously established (33) in 24-well plates with transwell inserts. Herein, inserts with 0.4 μm (Corning, #353095) and 3 μm pores (Corning, #353096) were tested. Albumin endocytosis and transcytosis was evaluated by addition of 50 μg/ml Alexa Fluor^™^ 594-albumin or Alexa Fluor^™^ 488-albumin to the top of the assay, and culture for 30 min. Endpoints included (1) endocytosis within endothelial cells, (2) albumin transcytosis to the lower fluid compartment, and (3) albumin transcytosis and uptake into astrocytes in the lower compartment (see Supplementary methods). For inhibitor studies, after establishment of BBB/BTB endothelial cells were serum deprived for 1 hour in Opti-MEM. Cells were pre-treated with inhibitors for 30 min at nontoxic concentrations (see Supplementary methods).

### shRNA-mediated knockdown of endothelial clathrin-independent endocytosis (CIE) genes

shRNA knockdown of genes associated with CIE was performed in bEnd.3 cells as described in Supplemental methods. Sequence of different shRNA lentiviral particle is given in Supplementary table 1.

### Glycoprotein Isolation

Glycoproteins were isolated using a glycoprotein isolation kit (89804; Thermo Fisher Scientific) as per the manufacturer’s protocol. 50 μg of proteins were loaded on SDS PAGE gel for assessing the level of FcRn and SPARC glycosylation using their antibodies.

### Co-immunoprecipitation (Co-IP)

Protein-protein interactions between Galectin-3 and FcRn were studied using Co-Immunoprecipitation Kit (ab206996, Abcam) (see Supplementary methods).

### Colocalization using confocal microscopy

Mouse bEnd.3 endothelial cells were cultured in chamber slides and stained by immunofluorescence for FcRn or Galectin-3 as described in Supplemental methods. Confocal microscopy was used to visualize staining.

### Immufluorescence (IF) of frozen human brain metastasis specimens

IF staining was performed as described previously on flash frozen human brain metastasis craniotomy specimens, collected under approved protocols (see Supplementary methods).

### Immunohistochemistry (IHC) of human brain metastasis specimens

Formalin-fixed, paraffin embedded blocks of human craniotomy specimens were collected as described in Supplementary methods. All procedures were performed according to the manufacturer’s instructions (LifeSpan, Inc.). The immunoreactivity was scored semi-quantitatively on a 0-3+ intensity basis in the endothelial, cancer, and brain parenchymal compartments.

### Graphic representation and statistical analysis

See Supplementary methods.

## Results

### Comparison of three transcytotic pathway ligands for brain metastatic distribution

Far-red emitting fluorescent labeled ligands/proteins for three transcytosis pathways in the CNS preclinical literature were compared *in vivo* to identify those with maximal penetration of brain metastases as compared to uninvolved brain. Tumor cells from two model systems, the HER2+ JIMT1-BR and triple negative MDA-MB-231-BR (231-BR) were each injected into the left ventricle of nude mice and metastases permitted to form. Both brain-tropic cell lines were EGFP labeled but expression was unstable *in vivo.* On the day of necropsy animals received an intravenous (IV) dose of far-red labeled transferrin or LRP1 peptide or Alexa Fluor^™^ 647-albumin, also far-red emitting. Based on an analysis of paracellular and transcytotic pathways in stroke (34), both a short (30m-1h) and a longer (4-6h) circulation times were used, before mice were perfused to remove probe from the bloodstream and necropsied (Fig. 1A). Whole brain sections were scanned and image analysis of probe intensity performed on 10 macro- and 10 micrometastases/brain that were detected by green fluorescence and confirmed by H&E staining of the adjacent tissue section. Ten areas of uninvolved brain/mouse were also imaged.

**Figure 1.**
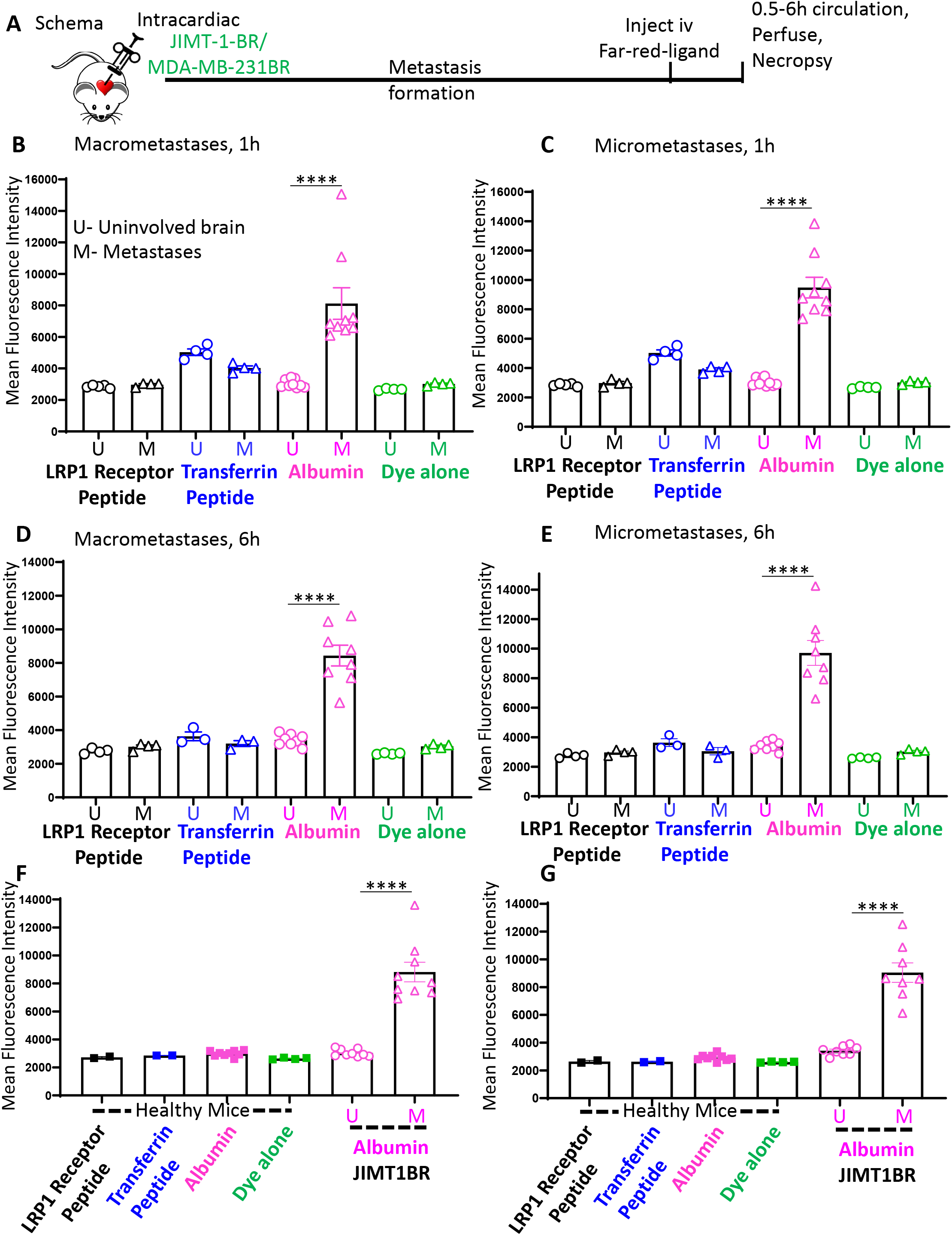
Comparison of three transcytosis pathways for uptake in the HER2+ JIMT1-BR experimental brain metastasis model of breast cancer. A. Experimental design. Mice were injected with enhanced green fluorescent protein (eGFP) labeled JIMT1-BR HER2+ brain-tropic breast tumor cells (JIMT1-BR) or MDA-MB-231-BR brain-tropic triple negative breast tumor cells (231-BR) into the left cardiac ventricle. Tumor cells circulated throughout the body and brain metastases developed in 3-10 weeks. Before necropsy mice were injected with Far-red emitting fluorescent labeled ligands/proteins (Transferrin receptor peptide, TfR; Low density lipoprotein receptor 1 peptide, LRP1; Albumin) which circulated for a short (0.5-1h) or longer (4-6h) period. Mice were perfused under deep anesthesia to remove label from the circulation. Whole brain sections were scanned for GFP and Far-red intensity in macrometastases and micrometastases. B-E. Mean and individual animal levels of each probe fluorescence intensity in uninvolved (metastasis free, U) brain and brain metastases (M), by metastasis size and time of circulation. F-G. The experiment was repeated with normal (uninjected) mice at two circulation timepoints, with albumin data from B-E shown for comparison. Statistical differences were calculated using One-way analysis of variance (ANOVA) by comparing uninvolved brain (U) to metastases (M) across all the groups. ****, P < 0.0001.

Graphs depicting ligand intensity for macrometastases and micrometastases in the JIMT1-BR brain-tropic model are shown on Fig. 1B–E; graphs combining the data for all metastases are on Supplementary Fig. S1; ligand distribution in healthy mice are plotted on Fig. 1F–G. Representative images of brain sections, macrometastases and micrometastases are shown on Fig. 2. Injection of the far-red dye alone as a control showed a low level of expression in both the uninvolved brain and metastases. LRP1 peptide was present at only low concentrations in all lesions at all timepoints. Intensity did not differ between uninvolved brain and lesions and was not significantly different from dye alone. TfR peptide exhibited a distinct pattern of uptake: TfR appeared higher overall than the far-red control at the early timepoint (compared to dye alone all metastases 1h, P=0.0467 and micrometastases 1h, P=0.0468). TfR distribution was numerically but not statistically higher in uninvolved brain than in metastases (no significance in macrometastasis, micrometastases, or all metastases 1h P=0.8337, 6h P=0.9982), a trend that is obvious on immunofluorescent micrographs (white arrows) outside of the metastatic lesions (Fig 2A–B). In contrast, Far-red-albumin demonstrated significantly greater uptake in metastases than uninvolved brain at all timepoints (compared to uninvolved brain-P<0.0001 for all metastases, macrometastases and micrometastases at 1h and 6hrs). Albumin uptake was similar in both micrometastatic and macrometastatic lesions, the latter comparable in length to a 3mm MRI-detectable lesion in a human brain. Albumin distribution within metastases was heterogeneous in intensity but widespread (Fig. 2A–B). Far-red-albumin levels in uninvolved brain were not significantly different from the dye alone control (All metastases 1h P=0.9988, 6h P=0.8764-Supplemenary Fig. S1A-B).

**Figure 2.**
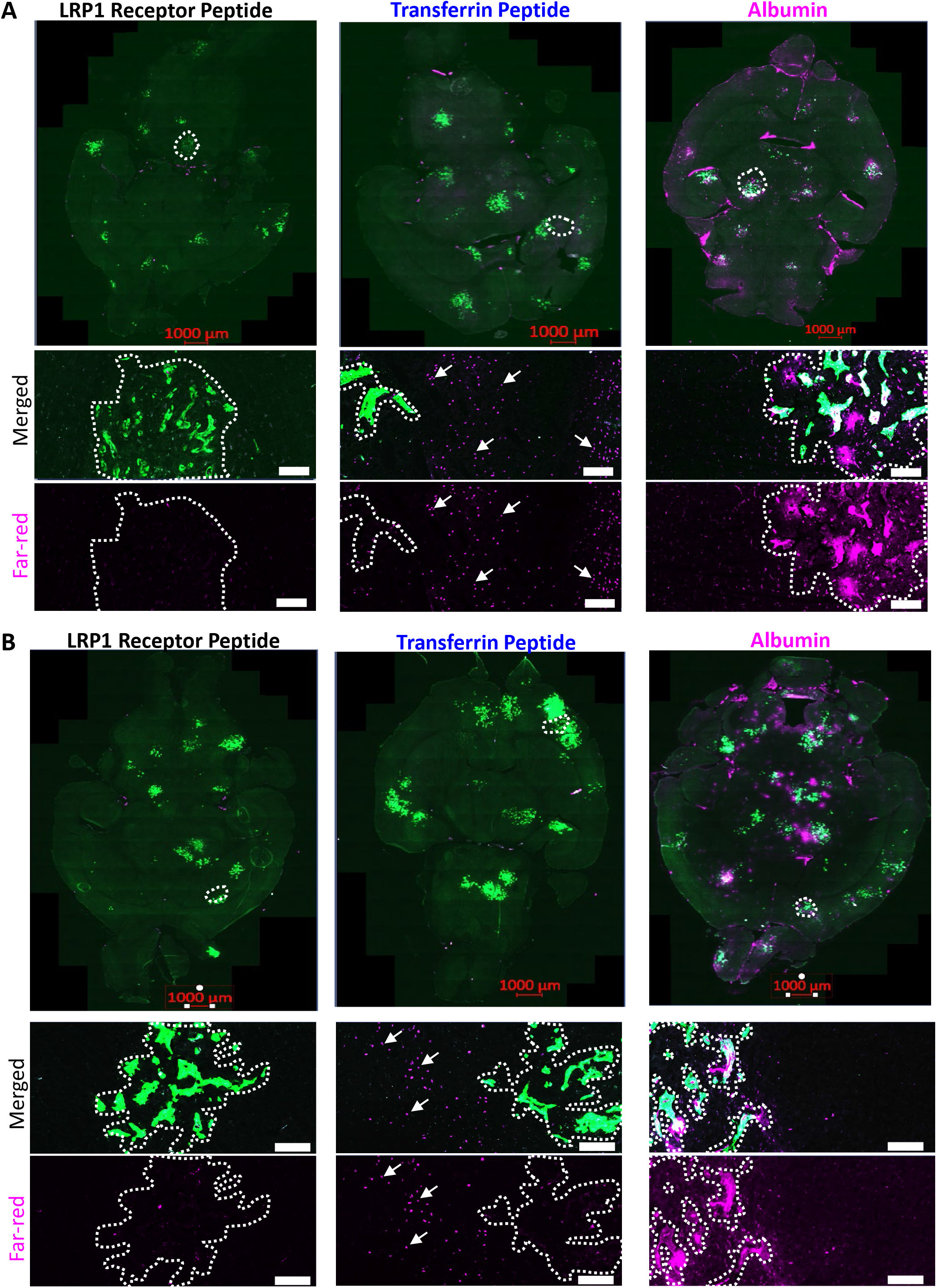
Photomicrographs of three transcytosis pathways for uptake in the HER2+ JIMT1-BR experimental brain metastasis model of breast cancer. For the experiment described in Fig. 1, representative photomicrographs of whole brain sections and metastases from each section (white dotted line) are shown for each transcytosis ligand and circulation time. For each metastasis a merged view of Far-red fluorescence of the transcytosis ligand and eGFP from tumor cells is shown and, beneath it, a Far-red channel only photo. Scale bar 125 μm. White arrows highlight TfR uptake in uninvolved brain.

In a separate experiment each probe was injected into healthy mice that had not received tumor cells (Fig. 1F–G). No significant differences were observed between dye alone and any of the transcytotic ligands at either timepoint. Data on albumin transcytosis at each timepoint is shown for comparison, suggesting that uptake in uninvolved brain was equivalent to normal brain.

Similar trends were apparent in the 231-BR brain-tropic triple negative metastasis model (Fig. 3A–F, Supplementary Figs S2-S3). LRP-1 peptide distribution into metastases was not significantly distinct from the dye alone control and difficult to visualize (Fig. 3E–F). TfR peptide distribution was higher in uninvolved brain than brain metastases at the 0.5-1h timepoint (compared to uninvolved dye alone-all metastases 1h P=0.0157, macrometastases P=0.0055 and micrometastases P=0.0935; uninvolved brain vs metastases-no significance), but near the far-red control at the longer 4-6h timepoint (no significance compared to dye alone uninvolved brain or uninvolved brain vs. metastases). Images confirmed a higher TfR distribution in uninvolved brain (white arrows) than in metastases (Fig.3E–F). Far-red-albumin distribution again demonstrated significant uptake into macrometastases and micrometastases at all timepoints, higher than that of uninvolved brain or dye control (Compared to uninvolved brain-P<0.0001 for all metastases, macrometastases and micrometastases at 0.5-1h and 4-6hrs). Albumin distribution was heterogeneous within a macrometastasis; occasional spots of albumin intensity in uninvolved brain often corresponded to single-few infiltrating tumor cells (Fig. 3E–F).

**Figure 3.**
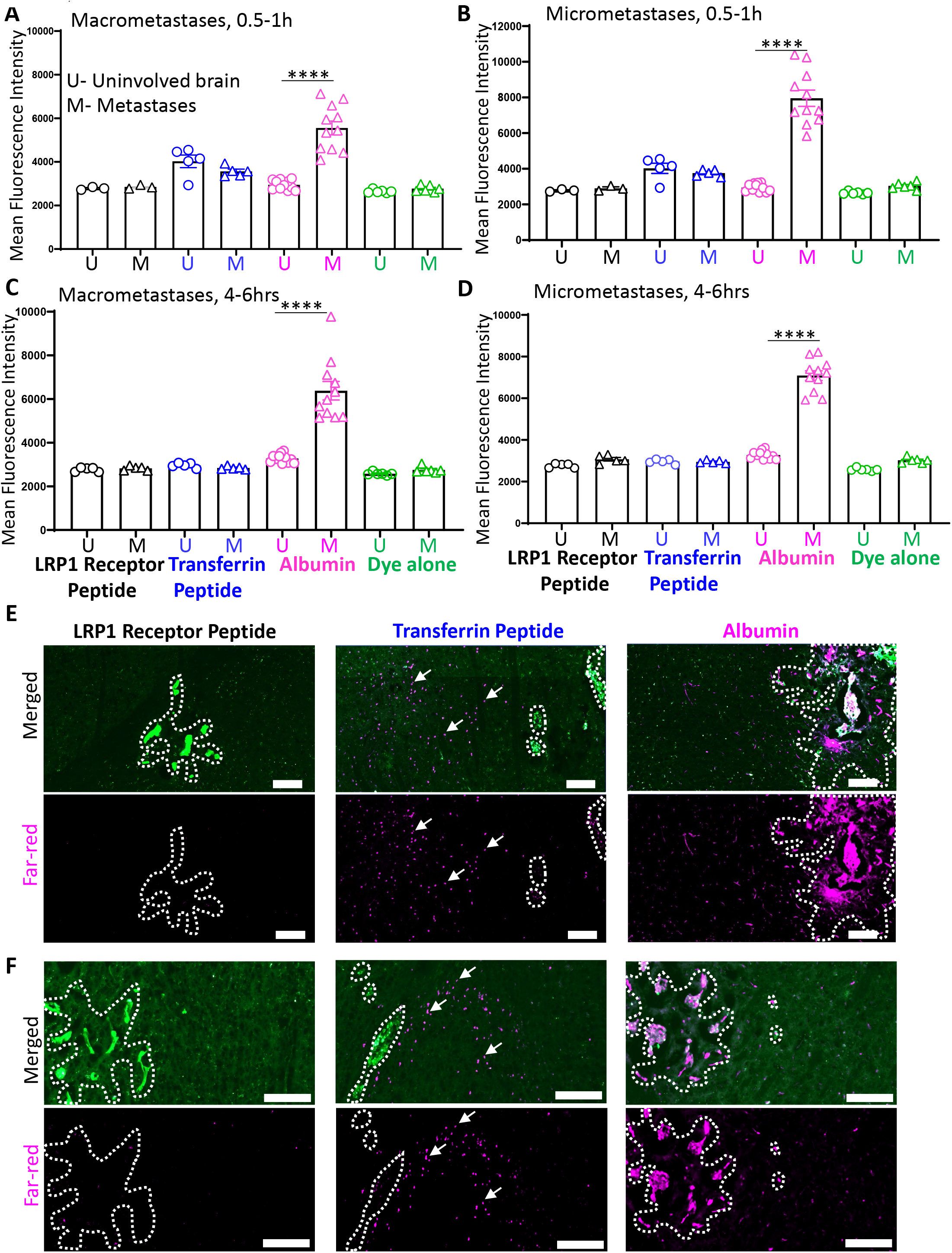
Comparison of three transcytosis pathways for uptake in the triple-negative 231-BR experimental brain metastasis model of breast cancer. An experiment described in Fig 1A was conducted using MDA-MB-231-BR (231-BR) brain-tropic cells. A-D. Mean and individual animal fluorescence intensity levels of each probe in uninvolved (metastasis free, U) brain and brain metastases (M), by metastasis size and time of circulation. E-F. Representative photomicrographs of metastases from each experimental arm (white dotted line) showing a merged view of Far-red fluorescence of the transcytosis ligand and eGFP from tumor cells along with far-red channel alone. Magnification scale bars as shown (125 μm). White arrows highlight TfR uptake in uninvolved brain. Statistical differences were calculated using One-way analysis of variance (ANOVA) by comparing uninvolved brain (U) to metastases (M) across all the groups. ****, P < 0.0001.

Taken together, the data establish that the three well-known transcytotic pathways studied are present at relatively low intensities in uninvolved brain. Only one of the three pathways, albumin, consistently and significantly distributed to brain metastases. Albumin uptake was comparable in macrometastases, the object of therapeutic experiments, as well as micrometastases that may be important to prevention of future metastatic outgrowth. Because of its promising distribution pattern, we studied albumin uptake into metastases further.

### Albumin transcytosis is distinct from paracellular marker distribution

We asked if albumin distribution into experimental metastases differed from that of a more typical paracellular entry method. Mice with 231-BR or JIMT1-BR brain metastases were dosed IV with Far-red-Albumin (Albumin-Alexa Fluor^™^ 647) and Biocytin TMR (Tetramethylrhodamine Biocytin), a compound of approximate drug size (869 Da) used as a probe of paracellular permeability (34,35), before perfusion and necropsy. Data for all metastases, separated into macro- and micrometastases are shown on Fig. 4A–B. Representative pictures showing distribution of Far-red-Albumin and Biocytin TMR in JIMT1-BR and 231-BR metastases are shown on Fig.4C and D, respectively. In the JIMT1-BR model system distribution of albumin and biocytin in uninvolved brain was relatively low and comparable at all timepoints (Fig. 4A). Biocytin expression showed a mean 73.2% and 68.5% increase in intensity as compared to uninvolved brain in macrometastases and micrometastases, respectively (macrometastases, P=0.0019; micrometastases, P=0.0006). This increased distribution was far less than that for albumin, which was 216.3% and 224.3% increased over uninvolved brain for macrometastases and micrometastases, respectively (P<0.0001 for both macrometastases and micrometastases). Similar trends were observed in the 231-BR model system, with albumin distribution a mean of 44.2% and 81.0% higher than uninvolved brain in macrometastases and micrometastases respectively (P<0.0001 for both macrometastases and micrometastases) (Fig. 4B). Photomicrographs show metastases in both model systems with observable albumin fluorescence and minimal if any discernable biocytin (Fig. 4C–D). The data show that albumin distribution to metastases in two model systems is distinct and generally higher than a paracellular permeation probe.

**Figure 4.**
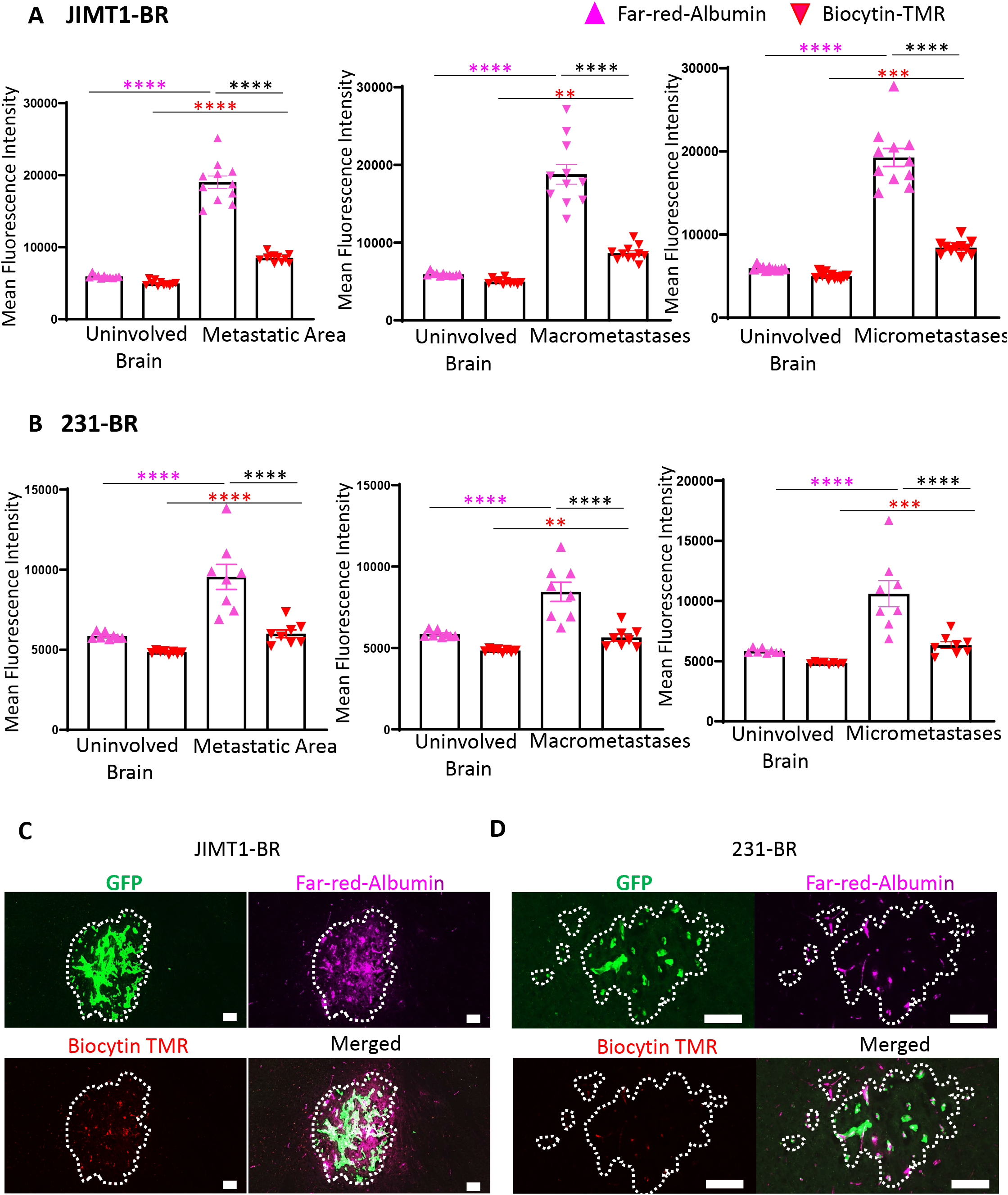
Far-red-Albumin distribution to metastatic colonies and micrometastases is distinct from paracellular distribution. Mice with either JIMT1-BR or 231-BR metastases received IV Far-red-Albumin and Tetramethylrhodamine Biocytin (Biocytin TMR), a marker of paracellular permeability; probes circulated for 60 and 10 min, respectively, followed by perfusion. All lesions in a whole brain section were analyzed for fluorescent uptake pattern. A-B. Means and individual animal data of each probe fluorescence intensity for uninvolved brain, all metastases (left), macrometastases (center) and micrometastases (right) are plotted. A. JIMT1-BR model system. B. 231-BR model system. C-D. Representative photomicrographs of a metastatic area with Far-red fluorescence of albumin, red fluorescence of Biocytin TMR, tumor cells expressing eGFP and their merge for JIMT1-BR (C) and 231-BR (D) model systems are shown. Scale bar 125 μm. Statistical differences were calculated using One-way analysis of variance (ANOVA) by comparing uninvolved brain to metastatic area/ macrometastases/ micrometastases across all the groups.

### Development of an *in vitro* model system incorporating albumin transcytosis

To ask what role the blood-brain barrier (BBB) and blood-tumor barrier (BTB) may be playing in these distribution patterns, we turned to *in vitro* models. We previously reported an *in vitro* model system for paracellular distribution of compounds through a BBB or BTB (33). Briefly, brain endothelial cells were cultured to confluence on the top side of a 0.4 μm porous filter, with pericytes cultured on the bottom side. The filter was then placed atop a chamber containing astrocytes in culture medium, with (BTB) or without (BBB) tumor cells (Fig. 5A). This model faithfully reflected Texas red dextran paracellular permeability *in vivo (36).* When Alexa Fluor^™^ 594-albumin was applied to this configuration, no endocytosis or transcytosis was observed. After many modifications, revision of one feature enabled transcytosis. Albumin endocytosis into endothelial cells (Fig. 5B) and transcytosis into the lower chamber (Fig. 5C) was observed only with wider, 3.0 μm pores. Permeability of doxorubicin via paracellular permeability was slightly higher in BTB than BBB cultures but did not vary with pore size (Fig. 5D). To ask how the larger pores could influence endocytosis, confocal microscopy was performed through the width of the 3.0 μm porous filter and demonstrated that cell processes from phalloidin/NG2 stained pericytes invaded through these larger pores to establish connections with the endothelium (Figure 5E), similar to their peg-and-socket physical interaction in the normal BBB (37). Transcytosis of albumin in this model was also confirmed by its presence on astrocytes in the lower chamber (Fig. 5F–G); both brain-tropic cell lines produced comparable results (Fig. 5F–H). Albumin costained with Rab11a, a vesicular trafficking protein known to be involved in transcytosis (38), in the endothelial layer (Fig. 5I).

**Figure 5.**
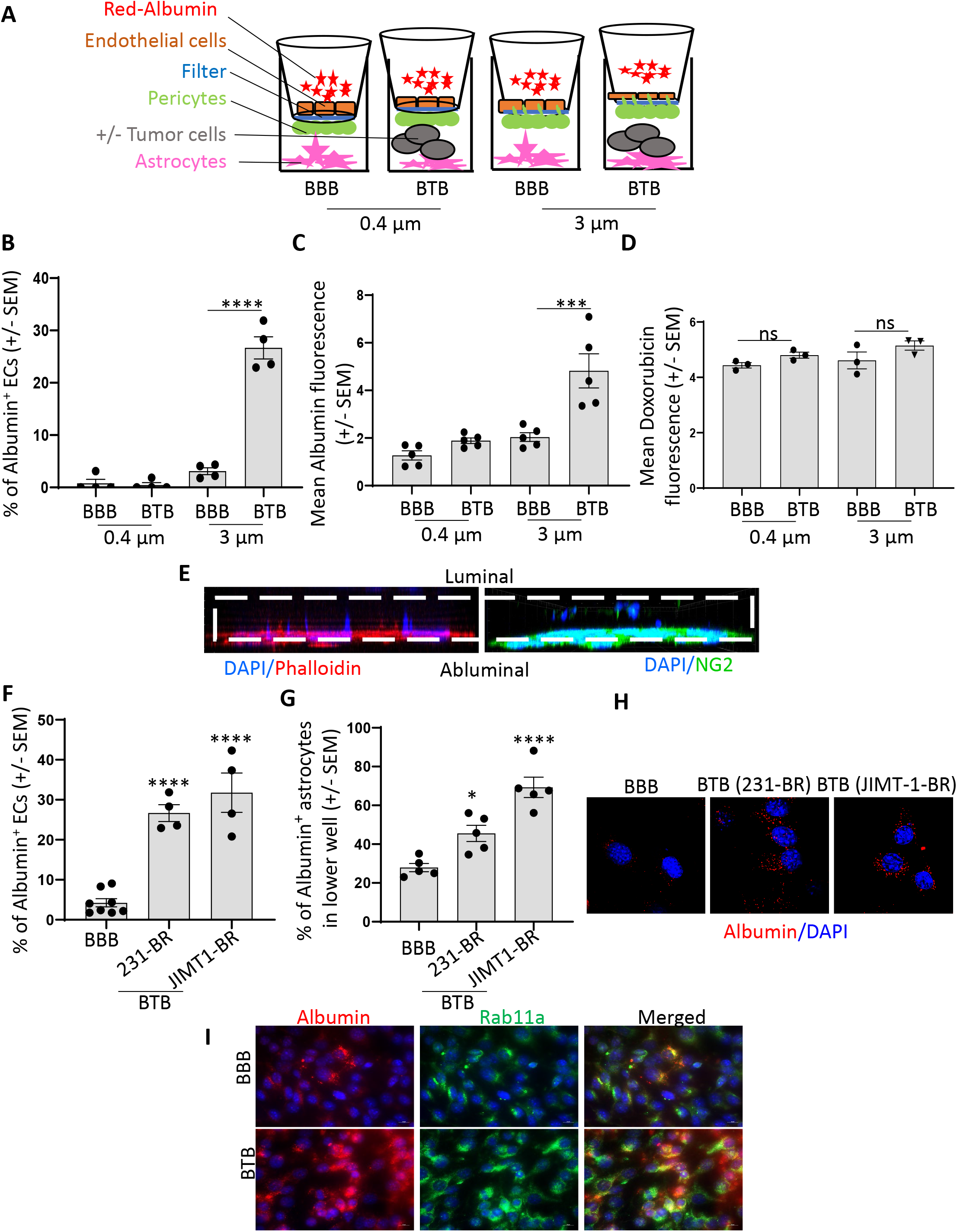
An *in vitro* assay of BBB and BTB function that demonstrates albumin endocytosis and transcytosis. A-D. Comparison of cultures using 3 μm and 0.4 μm porous filters. A. Experimental design. 50 μg/ml Alexa Fluor^™^ 594-albumin was applied to the top chamber and incubated for 30 min. B. Percentage of endothelial cells exhibiting albumin endocytosis. C. Transcytosis of Alexa Fluor^™^ 594-albumin after 2 hrs incubation was measured by Red fluorescence in the lower chamber (n=5). D. Application of doxorubicin to the *in vitro* assays for 45 min demonstrated no effect of filter pore size on paracellular permeability, quantified by doxorubicin fluorescence in the bottom well (n=3). E. Confocal microscopy through a 3 μm filter from a completed BTB assay stained for pericytes with NG2 and DAPI or Phalloidin and DAPI; demonstrating invasive NG2+ pericyte processes reaching through the larger pores toward endothelial cells. F-H. Data using 3 μm porous filters. F. Percentage of endothelial cells exhibiting albumin endocytosis in normal (blood-brain barrier, BBB) and tumor cell containing (blood-tumor barrier, BTB) cultures (n=4). G. Transcytosis of albumin through the cultures described in F onto astrocytes in the lower chamber (n=4). H. Photomicrographs of endothelial cells from F. Red, albumin; Blue, DAPI. I. Co-immunofluorescence of endothelial cells from BBB and BTB cultures for albumin (Red) and Rab11a (Green) with blue DAPI staining. Magnification bar, 10 μm. All experiments show mean +/- SEM with experiments conducted at least four times, until specified. Statistical differences were calculated using Oneway analysis of variance (ANOVA).

### Albumin transcytosis by macropinocytosis, but dependent on FcRn

To translate albumin uptake patterns into effective drug delivery, an understanding of its permeation patterns in the normal brain versus metastases is needed. Herein, we investigated the endocytic process by which albumin penetrates endothelia (Fig. 6A–B). Incubation of *in vitro* BTB cultures with Fillipin III or Chlorpromazine, inhibitors of caveolin-dependent (39) and clathrin-dependent (40) endocytosis, respectively, had no significant effect on albumin endocytosis. Ethylisopropylamiloride (EIPA), an inhibitor of macropinocytosis (41), inhibited albumin endothelial endocytosis by 95.5% (p<0.0001) in JIMT1-BR (Figure 6A) and 80.3% (p<0.0001) in 231-BR model systems (Figure 6B). Macropinocytosis, translated as “big drinking”, is the intake of fluid and soluble constituents in large vesicles. As a nonselective process, however, fluid and whatever is dissolved in it is taken up in classical macropinocytosis. It would make little sense that chemotherapeutic drugs would not be similarly taken up and readily distributed to brain metastases, in contrast to voluminous experimental data.

**Figure 6.**
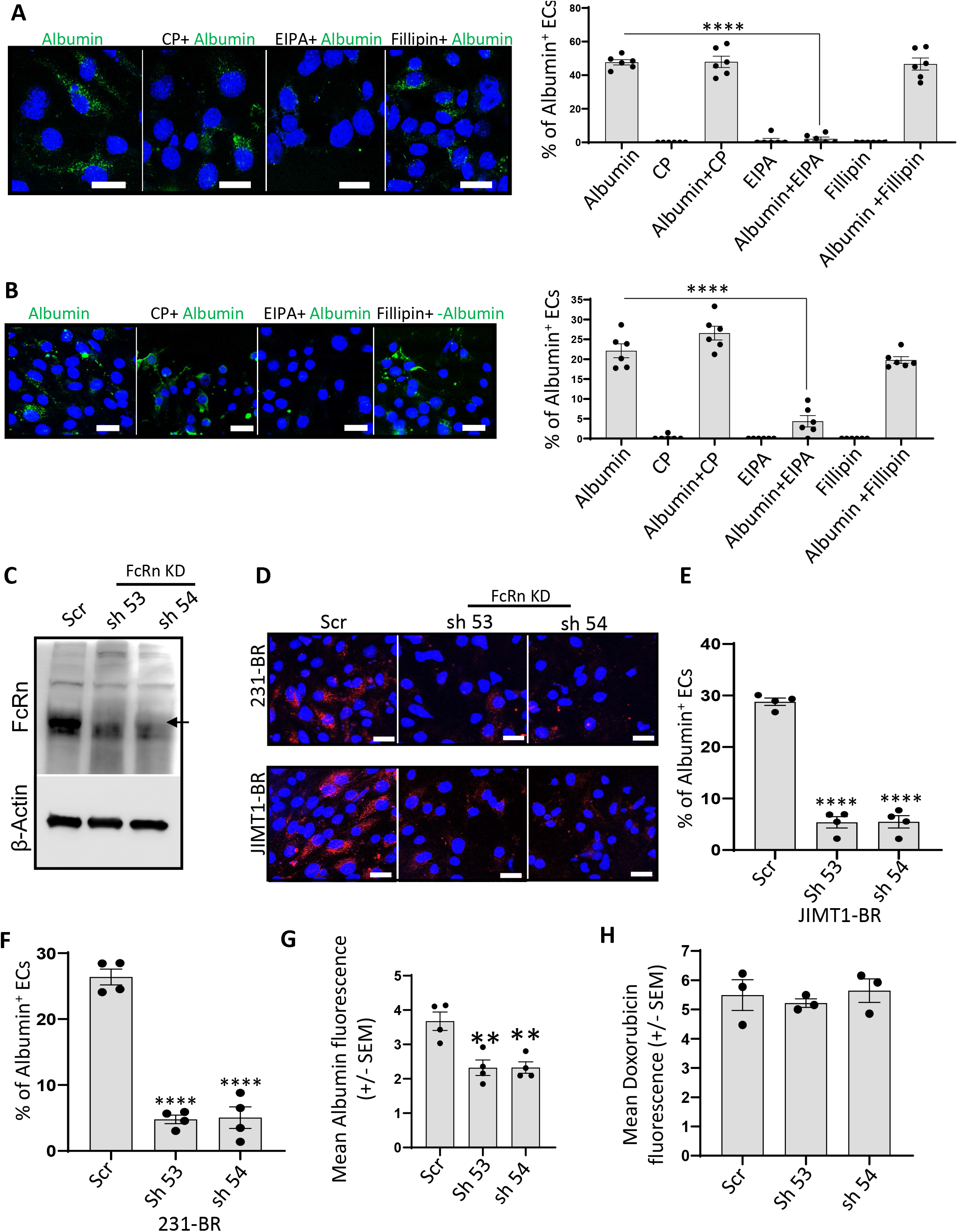
Albumin transcytosis through the blood-tumor barrier (BTB) uses macropinocytosis, but dependent on FcRn. A-B. Using the *in vitro* assay of BTB function (Fig. 5A), inhibitors of endocytic pathways were applied 30 min before addition of 50 μg/ml Alexa Fluor^™^ 488-albumin for 30 min. Brain endothelial endocytosis of albumin was quantified in cultures with: A. JIMT1-BR or B. 231-BR tumor cells, respectively (n=6). Scale bar 20 μm. C-H. Endothelial knockdown of FcRn disrupts albumin endocytosis and transcytosis. C. FcRn was knocked down in mouse brain endothelial cells using two independent shRNAs; the FcRn band is the top of a doublet as indicated by an arrow. D-G. Alexa Fluor^™^ 594-albumin was applied to *in vitro* BTB cultures and its endocytosis into endothelial cells was evaluated. D. Representative photomicrographs. Red, Albumin; Blue, DAPI. E-F. Percentage of albumin+ endothelial cells in JIMT1-BR and 231-BR BTB assays, respectively. G. Transcytosis assays measuring albumin fluorescence into the lower chamber (n=4). H. Paracellular movement of doxorubicin in 231-BR model BTB culture into culture medium of lower well.

A potential answer to this conundrum lies within the general category of macropinocytosis: A protein-binding macropinocytotic pathway has been described, termed clathrin-independent endocytosis (CIE), also known as clathrin-independent carrier (CLIC), glycolipid and lectin hypothesis (GL-Lec), or GPI-enriched compartment (GEEC). As an example, the membrane protein CD44, via extracellular N-glycosylation, binds galectin 3 (Gal-3) which, as a pentamer, coalesces multiple CD44 proteins. When coalesced in a membrane region containing high levels of glycosphingolipids, an endocytic pit forms (42). If this mechanism were to apply to albumin, it must bind an endothelial membrane protein. At least two prominent endothelial proteins have been reported to bind albumin, FcRn (27) and secreted protein acidic and rich in cysteine (SPARC or Osteonectin) (28). An albumin binding protein gp60 was previously reported but we could not identify it with current antibodies on western blots.

FcRn is highly expressed in normal BBB endothelia where it binds and recycles both albumin and IgG back to the bloodstream (43). Partial knockdown of FcRn in brain endothelial cells by two shRNAs was performed (Fig. 6C). When albumin was incubated with models expressing either shFcRn or scrambled shRNA, endocytosis was significantly reduced (>80%, p<0.0001) in the FcRn knockdowns (Fig. 6D); reduced albumin endocytosis was noted in BTB cultures containing either JIMT1-BR or 231-BR tumor cells (Fig. 6E–F). FcRn knockdown also reduced transcytosis of albumin to the lower chamber of BTB cultures by ~37% (P=0.004) (Fig. 6G). No effect on doxorubicin paracellular permeability was observed (Fig. 6H). The data indicate that a major portion of albumin endocytosis and transcytosis in the BTB is not via nonselective macropinocytosis but involved an FcRn-mediated pathway.

### Albumin macropinocytosis through the BTB displays features of CIE

We asked if albumin endocytosis, via FcRn binding, fit a general profile of CIE described for CD44 *in vitro.* Brain endothelial FcRn was N-glycosylated (Fig. 7A), in agreement with previous data in rat kidney cells (44). Brain endothelial FcRn bound Gal-3 in two-way co-immunoprecipitations (Fig. 7B–C, input controls on Supplementary Fig. S4). Gal-3 costained with FcRn in endothelial cultures using immunofluorescence experiments (Fig. 7D). Gal-3 was then knocked down from brain endothelial cells using two shRNAs (Fig. 7E). When these cells were incubated with albumin, endocytosis was reduced by 80% in the Gal-3 shRNA cultures (p<0.0001) (Figure 7F–H). A reduction in albumin endocytosis was also observed if membrane glycosphingolipids were depleted from endothelial cells using a Glucosylceramide synthase (GCS) inhibitor (45) (Figure 7I–K). Endocytosis of albumin was dynamin-independent (Supplementary Fig. S5A-D) and dependent on cdc42 (Supplementary Fig. S6A-B), also consistent with a CIE mechanism.

**Figure 7.**
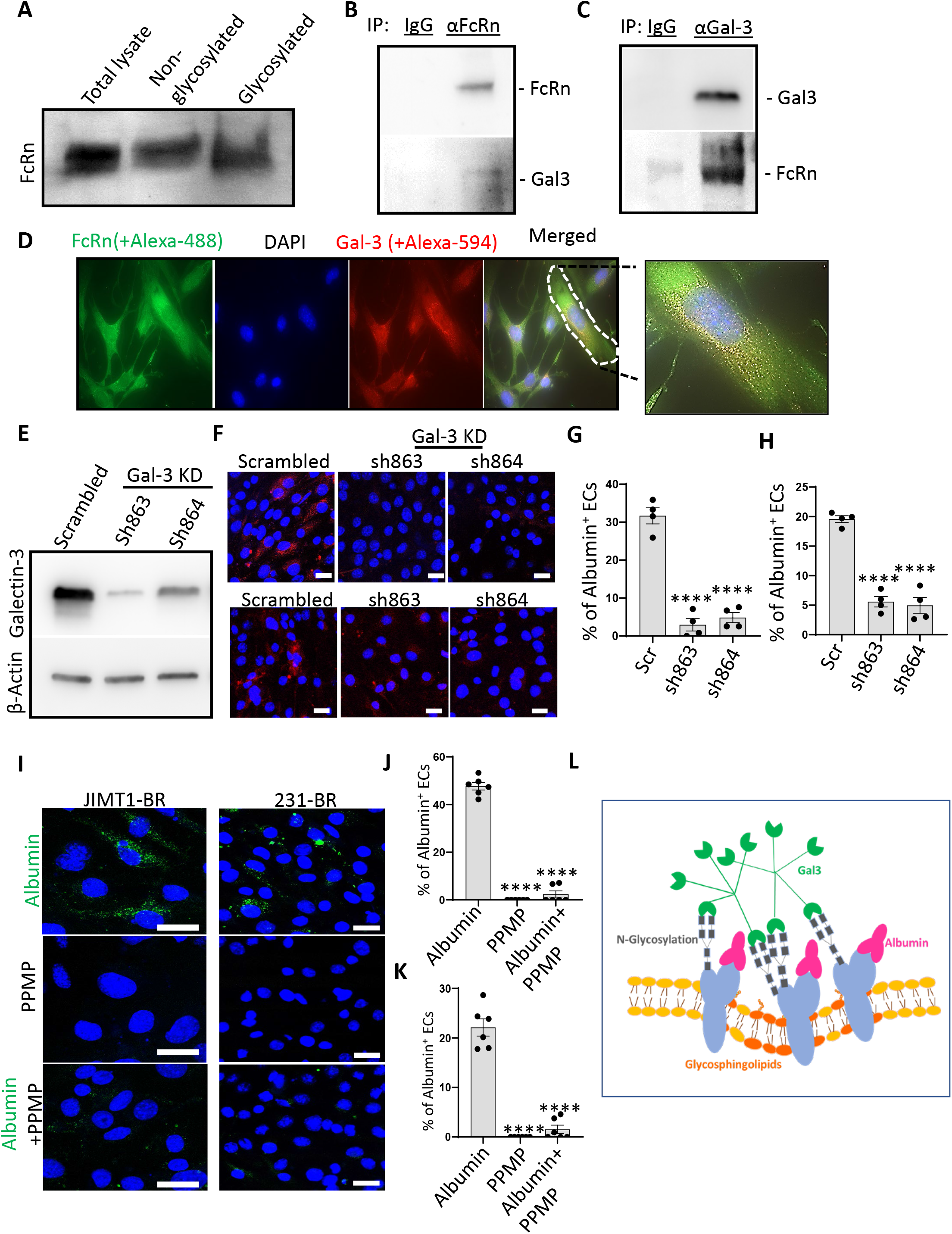
Albumin transcytosis through blood-tumor barrier (BTB) cultures demonstrates features of clathrin-independent endocytosis (CIE). A. A lysate of brain endothelial cells was separated into glycosylated and non-glycosylated fractions using Concanavilin A based glycoprotein isolation kit and probed for FcRn. B-C. Brain endothelial FcRn bound Galectin-3 (Gal-3) in two-way co-immunoprecipitations. D. Co-immunofluorescence of FcRn and Gal-3 in cultured brain endothelial cells. Green, FcRn; Red, Gal-3, Blue, DAPI. E. Western blot of Gal-3 knockdown by two independent shRNAs compared to a scrambled shRNA control in mice brain endothelial cells. F-H. Representative photographs (F) of endothelial endocytosis of Alexa Fluor^™^ 594-albumin using endothelial cells from **E** in an *in vitro* BTB assay on both model systems and its quantitation (G-JIMT1-BR, H-231-BR) (n=4); scale bar 10 μm. I-K. Glycosphingolipids are required for albumin endocytosis. Endothelial cells were treated with 30 μM of PPMP (DL-threo-1-Phenyl-2-palmitoylamino-3-morpholino-1-propanol) for 1 h to deplete glycosphingolipids. Endocytosis of Albumin was determined using an *in vitro* BTB assay with JIMT1-BR (I, J) and 231-BR (I, K) and representative photographs and quantitation are shown (n=6), scale bar 10 μm. All experiments were performed at least four times and statistical differences were calculated using One-way ANOVA. L. Schematic of albumin endocytosis into the BTB endothelium. Albumin (pink) binds FcRn (blue) in regions of high membrane glycosphingolipids (orange). N-glycosylation of FcRn (gray) binds to pentameric Gal-3 (green) resulting in coalescence to form a membrane invagination.

SPARC is an extracellular matrix modifying protein made by brain endothelial cells with FcRn binding properties (46). shRNA knockdown of SPARC only heterogeneously decreased albumin endocytosis in two clones (Supplementary Fig. S7A-B). SPARC was also non-glycosylated (Supplementary Fig. S7C). These data eliminated SPARC from further consideration as an albumin CIE pathway.

Taken together, *in vitro* data using BBB and BTB cultures demonstrate that albumin likely transcytoses the BTB, at least in part, by a CIE mechanism (Fig. 7L). Data suggests that albumin binds FcRn, which as an N-glycosylated protein binds Gal-3, resulting in receptor aggregation in a sphingolipid dense membrane region, resulting in invagination.

### Albumin CIE proteins in human brain metastases

Whether the albumin CIE transcytotic pathway is evident in human brain metastases is important for potential translational development. Initial studies were conducted with a limited number of frozen craniotomy specimens, suitable for IF. A specimen, representative of 4/9 examined, shows localization of collagen IV to identify BTB capillaries and co-localization of Gal-3. The adjacent section was stained for FcRn and also shows patchy co-localization, which could be consistent with its coalescence into membrane domains (Fig. 8A).

**Figure 8.**
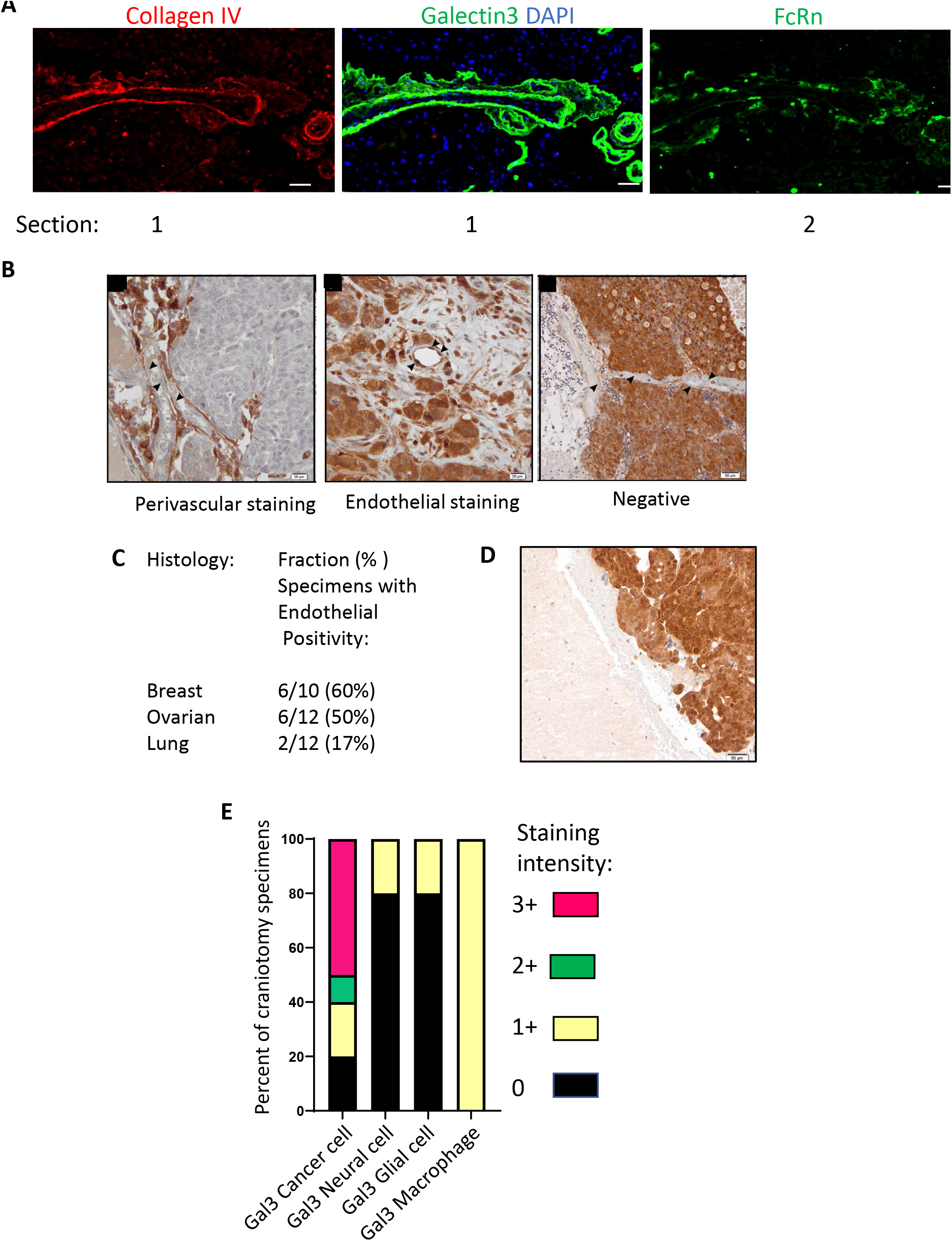
Galectin-3 (Gal-3) positivity is a consistent feature of the blood-tumor barrier of human brain metastases. A. Co-immunofluorescence of a snap frozen brain metastasis of human breast cancer showing Gal-3 colocalization with BTB vessels outlined by collagen IV. In an adjacent section the same structures demonstrate colocalization with FcRn. B-C. Gal-3 immunohistochemical staining of formalin-fixed, paraffin embedded resected human brain metastases. B. Representative photographs of Gal-3 staining patterns, on the luminal surface of, or within the BTB (black arrows highlighting perivascular, endothelial and negative staining-left to right). C. Percentage of craniotomy specimens from three cancer histologies expressing BTB endothelial Gal-3. D. Single example of a craniotomy specimen with a border of uninvolved brain, which demonstrated less Gal-3 staining. E. Gal-3 staining patterns of other components of brain metastases of breast cancer, expressed as a percentage of specimens with 0-3+ intensity staining (n=10). Scale bars: 50 μm.

Additional studies were performed by IHC using formalin-fixed, paraffin embedded craniotomy specimens from multiple cancer histologies. Staining for Gal-3 on metastatic capillary endothelial cells was found in 60% of breast cancer and 50% of ovarian cancer craniotomies but only 17% of lung cancer specimens (Fig. 8B–C). Staining was intracellular, which could reflect synthesis or intracellular sorting and trafficking, and luminal, consistent with an extracellular glycocalyx/CIE structure. A single specimen contained a margin of uninvolved brain in which Gal-3 was less prominent (Figure 8D). Interestingly, recent data comparing primary High grade serous ovarian cancer to its matched brain metastases highlights alteration of a complex network included the intracellular vesicular transport, along with cell cycle, lipid metabolism, cell junction organization etc. (47).

In addition to brain metastasis endothelial cells, Gal-3 expression was quantified on cancer cells, neurons, glial cells and macrophages in craniotomies on a 0-3+ intensity scale (Fig. 8E, Supplementary Fig. S8). While mainly absent on brain cells, intense staining was seen in cancer cells. These data suggest that a drug delivery via the CIE pathway could also operate in tumor cells, but spare most normal brain neurons.

## Discussion

In an effort to improve therapy for patients with brain metastases, or primary brain cancers, multiple investigators have reported preclinical studies using drugs linked to, or encapsulated in transcytotic pathways operative in the brain. Few of these have made it into the clinic or been FDA approved. These studies differ in at least four respects: the transcytotic pathway being drugged; the drug; its formulation/dose/schedule; the model system. Unfortunately, multiple variables make comparisons and conclusions difficult. Herein, we have conducted a survey of one variable, the transcytosis pathway, as a fundamental step toward rational design of new, efficacious therapeutics.

Three well-known transcytotic pathways, TfR, LRP1 and albumin were all labeled with a far-red fluorophore in order to compare their distribution patterns side-by-side. Peptides were used for TfR and LRP1 to enable far-red labeling. The probes were circulated in animals bearing two hematogenous models of breast cancer brain metastases, each for two time periods, before probe in the circulation was removed by perfusion and tissue uptake quantified. Each transcytosis pathway produced a distinct distribution pattern. In addition, brain metastatic distribution did not covary with uninvolved brain distribution. These data strongly support the use of metastatic models for probe and drug distribution translational studies. It will be of interest to quantitate the distribution of these probes in primary CNS tumors to determine whether these trends are BTB- or tumor histology specific.

Albumin showed a distinct distribution pattern with features thought to be optimal for therapeutic translation. Almost all lesions were positive for albumin in both model systems and at both timepoints tested, suggesting it can effectively deliver a drug to metastases. Both macrometastases and micrometastases labeled comparably, which are targets for therapy and prevention of outgrowth of additional lesions, respectively. Albumin uptake by uninvolved or normal brain was comparable to the far-red dye control and nearly invisible on photomicrographs. Drug uptake into normal brain may be irrelevant if the target is not expressed, for instance a mutated oncogene. Alternatively, chemo- and molecular therapeutics that target wild type proteins found in the brain, if taken up into normal brain, could cause profound toxicities. Albumin distribution was also greater than, and independent of biocytin-TMR, a paracellular permeability probe, suggesting that this pathway may yield distinct uptake patterns from traditional paracellularly distributed drugs. The only potential weakness in albumin distribution that we observed was its heterogeneous intensity within a lesion, so that not all tumor cells would receive equal exposure.

Albumin is known to enter normal brain endothelia via binding to the FcRn, and to then be endocytosed and recycled back to the circulation, or directed to the lysosome for breakdown to provide amino acids, never reaching the brain parenchyma in significant concentrations. Antibodies also bind FcRn, and this contributes to their relatively long half-lives (27). We have investigated how albumin permeates the BTB using an improved *in vitro* model system. Many *in vitro* models of the BBB and BTB have been reported, each with strengths and deficiencies (48). The current assay was modified from a previously reported culture system that demonstrated paracellular drug and probe distribution (36). Enlarging the pores separating endothelia from pericytes allowed cell connections and transcytosis. Albumin endocytosed into endothelial cells and transcytosed past the pericytes into culture medium and onto astrocytes in the lower chamber. Endocytosis was largely by macropinocytosis and was dependent on FcRn, based on shRNA knockdown experiments. *In vitro* assays with brain endothelial cells demonstrated that FcRn, via N-glycosylation bound Gal-3, and worked in concert with membrane glycosphingolipids. These data support a BTB endocytic process termed clathrin-independent endocytosis (CIE). Future research will determine how albumin-linked cargoes are routed post-endocytosis to a transcytotic endpoint. We observed increased co-immunofluorescence staining of albumin with the Rab11a, a marker of transcytosis, in BTB compared to BBB models. Endothelial cells with features of CIE (such as Gal-3 staining) are present in the BTB of human craniotomy specimens. Also, the methods of drug traversal through the remaining BTB and parenchymal cells and matrices between the endothelium and tumor cells must be identified. Interestingly, tumor cells were highly Gal-3+ and macrophages (microglia) were uniformly Gal-3+ at lower levels, suggesting that an albumin CIE pathway may be operative in other parts of the metastatic brain for drug delivery.

Albumin has been a focus of research for many years with varying conclusions. Albumin can coat nanoparticles to increase their solubility and adsorption, such as nab-paclitaxel, but differing conclusions have been drawn over the stability of this formulation (26,49). Stability of albumin binding to drug (directly or indirectly) would be required for a transcytotic function in the CNS. Albumin has a track record of stable conjugation to peptides or drugs to improve their half-life or preferential accumulation pattern, which has led to FDA approval of Levemir ^®^ (insulin detemir) and similar products for enhancing insulin bioavailability and distribution (50–52). Nanoparticles have been coated with albumin in many preclinical experiments for metastatic cancer. A recent formulation of an albumin stably coating a nanoparticle and containing a PI3Kγ inhibitor and paclitaxel, when administered with checkpoint therapy, shrank metastatic mammary tumors in a genetically engineered mouse model (26) and may represent an example of an efficacious stable formulation that can be tested for brain metastases. Two reports of stable incorporation of albumin into nanoparticles have produced promising results in brain cancers: A BSA-tetramethylindotricarbocyanide iodide (DIR) nanoparticle showed limited distribution in the normal mouse brain but a several-fold enhancement in brains containing intracerebrally inoculated glioma; *in vitro* uptake of albumin-DIR by endothelial cells was sensitive to depletion of membrane cholesterol but did not alter the expression of paracellular permeability proteins such as ZO-1 or Occludin (25), consistent with a transcytotic mechanism. Wan *et al.* prepared human serum albumin-phosphatidylcholine nanoparticles containing the selective EGFR and HER2 small molecule inhibitor lapatinib. In a 4T1-luc brain metastasis model system, lapatinib-loaded nanoparticles distributed to the metastatic brain better than to the normal brain, and to a metastatic brain 5.4-fold better than free drug after IV injection. Brain metastatic mouse survival increased 61% in response to the nanoparticle formulation over free drug (53). Taken together, the data urge reconsideration of stably linked albumin as a transcytotic mechanism for drug treatment of brain metastases and possibly other CNS cancers.

## Acknowledgements

This project has been funded in whole or in part with Federal funds from the National Cancer Institute, National Institutes of Health, under Contract No. HHSN261201500003I. The content of this publication does not necessarily reflect the views or policies of the Department of Health and Human Services, nor does mention of trade names, commercial products, or organizations imply endorsement by the U.S. Government. The human tissue samples from France were provided by AP-HM tumor bank AC-2013-1786, BB-0033-00097.

