## Supplementary Methods for "Comparison of Three Transcytotic Pathways for Distribution to Brain Metastases of Breast Cancer"

**Supplemental Materials and Methods**

**Chemical Synthesis (General Experimental description)**

Fluorescent probes were prepared by the Chemistry and Synthesis Center, NHBLI, NIH, or purchased. Either Cy5 or Alexa Fluor™ 647, a sulfonated form of Cy5 with identical spectroscopic characteristics (Ex/Em wavelength: 650 nm/670-680 nm), was used as the fluorescent label. Adducts of either fluorophore are referred to with the prefix far-red.

Albumin from Bovine Serum (BSA), Alexa Fluor™ 647 conjugate (ref# A34785, **Far-red-Albumin**), 5-(and-6)-Tetramethylrhodamine Biocytin (**Biocytin TMR**, cat# T12921) were purchased from ThermoFisher Scientific. Transferrin and Angiopep (LRP1 receptor) peptides were synthesized in appropriately protected forms and coupled to Alexa Fluor™ 647 NHS ester (ThermoFisher Scientific, cat#A37566) (**Far-red-Transferrin peptide** and **Far-red-LRP1 Receptor peptide,** respectively) (1).

Reagents were purchased from commercial sources and used as received. Peptides were synthesized using an automated AAPPTEC Focus-XC-6V instrument running AAPPTEC software version 3.02.03. The sequences were assembled starting from the appropriately preloaded resin. Standard Fmoc chemistry was used to construct the required sequences. For coupling, 6.0 eq. of suitably protected amino acids (0.4M in DMF) were activated with HBTU (0.4M in DMF, 6.0 eq.) and NMM (0.8M, 12.0 eq.) and transferred to the peptide vessels under positive nitrogen pressure. After a coupling time of 60 min, the reagents were removed, and resins were washed with DMF. Fmoc- protecting groups were removed with 20% piperidine in DMF, the resin was washed with DMF and then taken to next coupling with the respective amino acid.

Preparative HPLC was performed on a 30.0 x 150 mm Waters XBridge® C18 column; particle size 5.0 microns with an elution rate of 30 mL/min and detection at 220 nm. Solvent A: water with 0.05% TFA; solvent B: acetonitrile with 0.05% TFA. Major peaks were collected and analyzed by LCMS. LCMS analysis was performed on an Agilent 1200 series HPLC system with an LC/MSC Trap XCT detector and a Zorbax 300SB-C18 3.5 um column (4.6 x 50 mm).

**Far-red-Transferrin peptide (**Transferrin peptide-Alexa Fluor™ 647 conjugate**)** (Ac-NH-Thr-His-Arg-Pro-Pro-Met-Ser-Pro-Val-Trp-Pro-CONHCH_2_CH_2_NH-Alexa647): Protected peptide AcNH-Thr(t-Bu)-His(Boc)-Arg(pbf)-Pro-Pro-Met-Ser(t-Bu)-Pro-Val-Trp(Boc)-Pro-CONHCH_2_CH_2_NH_2_ was synthesized according to the general method using 1, 2-diaminoethane trityl resin. After peptide synthesis, the terminal Fmoc was removed manually by incubating the resin with 20% piperidine for 15 min, followed by washing with DCM. The N-terminal amine was then acetylated by adding 10 mL pyridine and 7 mL acetic anhydride to the resin and shaking for 1 h. The resin was filtered off and washed with DCM 3 times. The protected peptide was then cleaved from the resin by treating 5x with a cocktail of 99:1:0.1 DCM:TFA:TIPSH for 15 min each, and the combined filtrates were evaporated under reduced pressure. The crude product was purified by RP flash chromatography (CombiFlash EZ, Teledyne ISCO, Inc) on a C18 column eluted with acetonitrile and water. The product fraction was freeze-dried to give a white powder. A mixture of the protected peptide (51 mg, 0.026 mmol), Alexa Fluor 647-NHS (25 mg, 0.020 mmol) and DIPEA (4 μL, 0.023 mmol) in anhydrous DMF (0.5 mL) was shaken at room temperature and monitored by HPLC. After the reaction was completed in 24 h, the solvent was evaporated under reduced pressure on a V10 apparatus (Biotage, Inc). The residue was incubated in 5 mL TFA cocktail (TFA/Phenol/TIPS/H_2_O 95/2/2/1) at room temperature for 1 h, and then the solvents were evaporated on the V10 apparatus. The residue was neutralized to pH 7-8 with aqueous saturated NaHCO_3_ solution (5 mL) and then purified by preparative HPLC with a gradient of 20 -> 40 % B over 10 min (Extended Data Fig. 1). A dark green solid product was obtained after freeze-drying the product fraction (25 mg, 74 % in yield), which was stored at 4°C prior to use. HPLC purity >95% at 647 nm; >84% at 280 nm. MS: Calcd for C_100_H_143_N_21_O_27_S_5_ (M+2H)/2: 1115.8; found 1115.8.

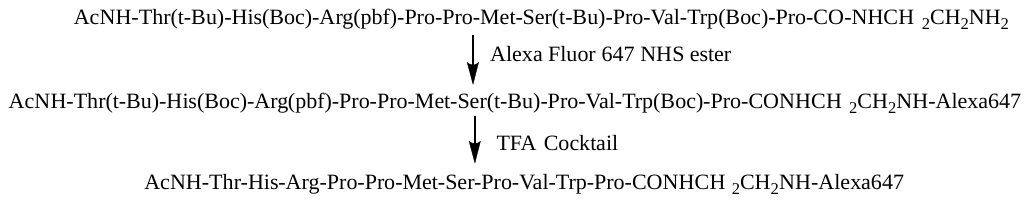

**Far-red-LRP1 Receptor peptide (**Angiopep-Alexa Fluor™ 647**)** (Ac-TFFYGGSRGK (Alexa647)RNNFK(Ac)TEEY-NH_2_): The peptide was synthesized according to the general procedure using Rink amide resin. Standard acid-labile side chain protecting groups (Pbf, Boc, *tert*-butyl) were employed with the exception of K15, which was installed with a Dde protecting group. Upon completion of the peptide sequence, the fully protected peptide on the resin was rinsed with DMF and transferred to a reaction vessel. The Dde protecting group was removed by shaking with 2% hydrazine in DMF (3 x 30 min at RT); the resin was rinsed 3 x with DMF, then treated with 20% piperidine in DMF (3 x 20 min at RT) followed by rinsing (3 x DMF). The partially deprotected material was acetylated at the N-terminus and K15 by shaking for 60 min at RT with 10 mL acetic anhydride, 10 mL pyridine and 2 g HOBt. After washing (3 x DMF) the peptide was deprotected and cleaved from the resin by treatment with 95:2.5:2.5 TFA: water: tri-isopropyl silane (30 min, RT). The filtrates were collected; the resin was washed 2x with the cleavage cocktail, and the volatiles were removed under reduced pressure. The residue was triturated with anhydrous ether and the product was isolated by preparative HPLC (gradient 15->40% B over 12 min) followed by lyophilization. The resulting peptide was conjugated with Alexa Fluor™ 647 following the procedure described above and the reaction was monitored by LCMS. Upon completion, the labeled peptide was isolated by preparative HPLC 15->40% B over 12 min, followed by lyophilization (Extended Data Fig. 2). HPLC purity: >98% at 220 nm. MS: Calcd for C_144_H_198_N_32_O_45_S_4_ (M+2H)/2 1613.8; found 1613.4.

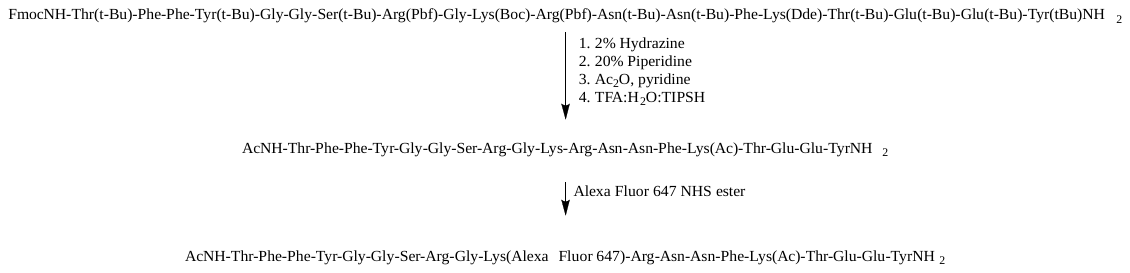

**Cell line origins and authentication****.** Two brain-tropic breast cancer cell lines, developed in the laboratory by serial passages of intracardiac injection, and harvesting and culturing of brain colonizing tumor cells, were utilized: the triple-negative 231-BR (2) and the HER2-overexpressing JIMT1-BR (3). Both lines expressed eGFP (enhanced green fluorescent protein) to aid in microscopic visualization. The 231-BR and the JIMT-1-BR lines were cultured in high Glucose DMEM (Dulbecco's Modified Eagle Medium, Gibco), with 10% Fetal Bovine serum and supplemented with 2mM L-glutamine for the JIMT1-BR. The JIMT1, MDA-MB-231 and -BR derivative lines have been analyzed by ATCC® using seventeen short tandem repeat (STR) loci and the gender determining locus. Only the MDA-MB-231 parental could be authenticated with the database of ATCC. The three brain-tropic variants matched their corresponding parental line.

Immortalized human astrocytes (HAL) and pericytes (HPL) were generated using SV40 Large T antigen as described previously (4). HAL were cultured in Astrocyte Medium (#1801, ScienCell) containing fetal bovine serum (FBS, Cat. No. 0010), astrocyte growth supplement (AGS, Cat. No. 1852) and penicillin/streptomycin solution (P/S, Cat. No. 0503) as per the manufacturer’s protocol. HPL were grown in Pericyte Medium (#1201, ScienCell) containing fetal bovine serum, pericyte growth supplement (PGS, Cat. No. 1252), and penicillin/streptomycin solution. The mouse brain endothelial cell line (bEnd.3) was cultured in Complete classic medium containing serum and Culture Boost (#4Z0-500, Cell Systems).

**Animal Experiments- General Information**. All animal experiments were performed under the regulation of the Animal Care and Use Committee (ACUC) of the National Cancer Institute (NCI), Frederick National Lab. NCI-Frederick is accredited by AAALAC International and follows the Public Health Service Policy for the Care and Use of Laboratory Animals. Animal care was provided in accordance with the procedures outlined in the “Guide for Care and Use of Laboratory Animals (National Research Council; 1996; National Academy Press; Washington, D.C.). The cell lines tested negative for a panel of viruses and for mycoplasma. Five-seven-week-old female athymic NIH nu/nu mice (Charles River Laboratories) were anesthetized with isoflurane/O_2_ and tumor cells were injected in the left cardiac ventricle each in 0.1 mL Ca^2+^ Mg^2+^ free PBS containing 1.75 x 10^5^ 231-BR or JIMT1-BR cells as previously described (3). Metastasis formation occurred over approximately 4 and 3 weeks, respectively. Animal health was monitored daily, and weights determined bi-weekly. Mice were euthanized individually before the preset endpoint when they displayed morbid symptoms, such as paralysis, head tilt, or loss of more than 20% of the body weight. At necropsy, brains were dissected, and hemispheres were fixed per the requirements of individual experiments. Sections were stored at −80 °C.

**Comparison of three transcytosis ligands.** For analysis of transcytosis, animals were randomized using StudyLog computer software. Each experimental arm consisted of ten animals injected per transcytosis ligand. After the development of brain metastases and on the day of necropsy, the fluorescent probes were solubilized in Dulbecco’s phosphate-buffered saline (D-PBS) with Ca^++^/Mg^++^. The probes were injected IV into the tail vein at 1% w/v (1mg in 100μl), following the protocol of Knowland *et al.*(5). The fluorescent probes circulated for 0.5–1h, or 4-6h to compare short and longer circulation times, respectively. Subsequently, the mice were perfused with Krebs-Ringer Bicarbonate buffer for one minute (pump rate at 10ml per minute) to eliminate the probes from the vasculature per previous protocol (4).

Relative molar equivalencies are: The transferrin peptide and LRP1 Receptor Peptide were injected at 30 and 21 times the molar equivalent of Albumin, respectively, using the weight equivalent of 1mg/mouse. The fluorescent marker Far-red-Dye alone (Alexa Fluor™ 647 NHS Ester (Succinimidyl Ester)) was injected using molar equivalent of Albumin- Alexa Fluor™ 647 conjugate (Molecular weight ~68,000Da, injected at 1mg/mouse, i.e., 0.0147 µmol/mouse).

At necropsy, brain and four systemic vital organs (liver, lungs, heart, kidneys) were dissected, fixed in 4% PFA for 4 hours at 4^○^C, immersed in 20% sucrose for 24 hrs, embedded in OCT and frozen in an ethanol/dry ice bath. This fixing method allowed the direct visualization the fluorescent probe. In several animals a brain hemisphere was directly frozen in OCT, enabling better tissue preservation for immunostaining. Brains were sectioned as follows: 5 series of three 8-µm-slices every 600µm, in sagittal sections. For the liver, lungs, heart and kidney, 2 series of three 8-µm-slices, 600µm apart were sectioned. The first slide in each series was stained with hematoxylin and eosin (H&E). The other sections were used for Whole brain sections scanning using Zeiss AxioScan.Z1 Slide Scanner for far-red and eGFP fluorescence. For quantitation analysis, whole brain sections were scanned for fluorescence (488/594/647) using Zeiss AxioScan.Z1 Slide Scanner. The adjacent H&E stained sections were also scanned using Zeiss AxioScan.Z1 Slide Scanner. For one section/mouse, located approximately 2400 µm deep from the lateral edge, all metastases- macrometastases (≥300 µm in length or 5000 µm^2^ area) and micrometastases (<300 µm in length or 5000 µm^2^ area) were localized/matched with H&E stained sections. Image analysis of probe intensity were performed on 10 macro- and 10 micrometastases/brain that were detected by green fluorescence and confirmed by H&E staining of the adjacent tissue section. Ten areas of uninvolved brain/mouse were also analyzed by ZEN 2 software. Mean Fluorescence Intensity values were used for plotting the distribution of all three transcytosis ligands in uninvolved area and metastases, along with dye alone control.

**Comparison of transcytosis versus paracellular permeability:** The above protocol was used to inject 231-BR and JIMT1-BR cells into mice and develop brain metastases. After brain metastasis formation and on the day of necropsy 100 µl of 1% Far-red-Albumin (Alexa Fluor™ 647-albumin) (Thermo Fisher Scientific # A34785) was injected into the tail vein (IV) of animals. Albumin was allowed to circulate for 1 h (total). After albumin has been circulating for 50 min, 1% (w/v) Biocytin-TMR (Thermo Fisher Scientific # T12921l) was injected (IV) and was allowed to circulate for 10 min, followed by the perfusion of animals as described above. Brains were harvested at necropsy and were sectioned as described above. This experiment included 11 animals/arm and was performed once.

For analysis, whole brain sections were scanned for fluorescence (488/594/647) using Zeiss AxioScan.Z1 Slide Scanner. The adjacent H&E stained sections were also scanned using Zeiss AxioScan.Z1 Slide Scanner. For one section/mouse, located approximately 2400 µm deep from the lateral edge, all metastases- macrometastases and micrometastases were localized on H&E stained sections. Image analysis of probe intensity were performed on 10 macro- and 10 micrometastases/brain that were detected by green fluorescence and confirmed by H&E staining of the adjacent tissue section. Ten areas of uninvolved brain/mouse were also analyzed by ZEN 2 software. Like above, Mean Fluorescence Intensity of far-red-albumin channel (647 nm) and biocytin-TMR in the red channel (594 nm), values were used for plotting the distribution of all three transcytosis ligands in uninvolved area and metastases, along with dye alone control.

***In vitro* blood-brain barrier (BBB) and blood-tumor barrier (BTB) assays.** The *in vitro* BBB/BTB assay reflecting paracellular permeability was established as previously described (4) in 24-well plates with transwell inserts. The generation and culture of immortalized (using SV40 Large T antigen) cell lines of pericytes and astrocytes were previously described (4). Inserts of 0.4 µm (Corning, #353095) and 3 µm pores (Corning, #353096) were tested herein to evaluate the best condition mirroring *in vivo* transcytosis. Briefly, immortalized human pericytes (HPLs) (3 × 10^4^), were seeded on abluminal side of transwell insert. Two hours later, immortalized mouse endothelial cells (bEnd.3) (1 × 10^5^) were seeded on the luminal side. The bottom of the 24-well was seeded with 1 × 10^5^ immortalized human astrocytes (HAL) in a different plate. To establish the BTB, 3-4 h later, 1 × 10^5^ cancer cells (231-BR or JIMT1-BR) were added to the astrocytes. The cultures were maintained for 24 hours (day 2). Subsequently, the inserts were placed onto the wells containing the astrocytes with or without the cancer cells and were maintained in pericyte medium (bottom chamber). The wells were incubated for 48 h to establish barrier features (day 4). On day 4, cells were serum starved for 1-2 hours in Opti-MEM. Transcytosis was evaluated by adding 50 μg/ml (500 µl in Opti-MEM) of Alexa Fluor™ 594-albumin (#A13101, Thermo Fisher Scientific) or Alexa Fluor™ 488-albumin (#A13100, Thermo Fisher Scientific) to the top of the insert (luminal side) and incubated for 30 min with three different readouts: (1) visualization of Alexa Fluor™ 594/488-albumin within the endothelial cells (endocytosis). The inserts were washed with cold PBS, fixed with 4% PFA and incubated with DAPI (1:500) to detect cell nuclei. Multiple random fields of the insert, with at least 10 cells per field, were photographed using confocal microscopy (Zeiss LSM 780). The number of endothelial cells with and without fluorescent albumin uptake were counted, then graphed as percent of albumin positive cells. (2) For Spectrophotometric measurement of Red-Albumin (Albumin-Alexa Fluor^TM^ 594) fluorescence in the lower compartment (transcytosis), 100 μg/ml (500 µl per well) of Red-Albumin was added to the upper culture and incubated for 30 min. Inserts were washed (both top and bottom) with PBS (3 times) and were placed in a 24-well plate containing 1ml PBS. Albumin endocytosed by the endothelial cells were allowed to transcytose to the lower chamber containing PBS for 60 or 120 min. Transcytosed albumin was measured by taking an aliquot of the lower chamber into a new plate (one reading per insert at two time points) and fluorescence was quantified using a plate reader (Ex- 590/Em- 622nm). (3) Visualizing Far-red Albumin (Albumin-Alexa Fluor^TM^ 647) in astrocytes (transcytosis): Like above, Far-red Albumin (100 μg/ml, 500 µl) was added to the upper culture and incubated for 30 min. Inserts were washed (both top and bottom) with PBS (3 times) and were placed in a 24-well plate containing 80% confluent astrocytes on coverslips. Albumin endocytosed by the endothelial cells were allowed to transcytose to the lower chamber for 60 min which was taken up by the astrocytes. Astrocytes were fixed with 4% PFA and incubated with DAPI to detect cell nuclei. Transcytosed albumin was counted as number of albumin positive astrocytes (albumin dots) under the fluorescent Zeiss microscope and quantified in five random fields per slide, each containing at least 10 cells.

To quantify paracellular permeability, the transwell inserts with endothelial cells and pericytes were placed in a new 24-well plate containing 1 ml PBS. Endothelial media containing 100 µg/ml doxorubicin (Sigma-Aldrich #44583) (Ex/Em:480/590 nm) was added on top of endothelial cells on the luminal side. 100 µl PBS was collected after 45 min incubation from 96-well clear bottom plates (Thermo Scientific, #165305). Fluorescence (at 480/590 nm) was measured in SpectraMax M2 (Molecular Devices). PBS was used as negative control. Statistical significance was determined by one-way ANOVA (nonparametric) using Turkey’s multiple comparison test

**Staining of pericytes.**

Once the BTB is established (day 4), the inserts were washed with PBS and cells were fixed with 4% PFA at room temperature for 15 min. The cells were blocked with 5% normal goat serum (Dako, # X090710-8) in PBS containing 0.2% Triton-X 100 for permeabilization. For pericytes, 1:100 diluted antibodies [ NG2 (Millipore, #AB5320)] was added in the lower wells and incubated overnight at 4 °C. Species-specific secondary antibodies (1:500 diluted in blocking reagent along with DAPI for nuclei counterstaining) were added and incubated in dark at room temperature for 45 min. A parallel BTB insert with pericytes alone in the abluminal side was also stained with DAPI and phalloidin. At the end, inserts were cut using a scalpel, placed on a glass slide, mounted with fluorescent mounting medium (Dako, # S3023) and imaged using confocal microscopy.

**Transcytosis inhibitor treatments**. For inhibitors studies, after establishment of BBB/BTB (on day 4) endothelial cells were serum deprived for 1 hr in Opti-MEM. Cells were pre-treated with inhibitors for 30 min [CP (Chlorpromazine)- 50 µg/ml, #C8138, Sigma Aldrich; Filipin III- 4 µg/ml, #F4767, Sigma Aldrich; EIPA/Amiloride -75 µM, #A3085, Sigma Aldrich; DL-PPMP- 30 µM, 60 min pretreatment, #BMLSL2150025, Enzo Life Sciences; MiTMAB- 2µM, #ab120466, Abcam). Alexa Fluor™ 488/594-albumin was added (50 μg/ml) to the top well (luminal side) and incubated for 30 min with or without inhibitors. Alexa Fluor™ 594/488-albumin uptake in the endothelial cells was visualized as per the above protocol using confocal microscopy. Statistical significance was calculated by one-way ANOVA (nonparametric) using Turkey’s multiple comparison test.

**shRNA-mediated knockdown of endothelial clathrin-independent endocytosis (CIE) genes.** The mouse brain endothelial cell (bEnd.3) line was cultured in Complete classic medium with serum and Culture Boost (#4Z0-500, Cell Systems). shRNA mediated knockdown of different genes was performed using MISSION® shRNA, lentiviral particles (Millipore Sigma, USA). Sequence of different shRNA lentiviral particle is given in Supplementary Table 1. Mouse brain endothelial cell (5x10^4^) were plated in 24 well plate and were allowed to grow for 24 h. Cells were transduced with 50 µl of lentiviral particle (Per reaction: 50 µl lentivirus+ 400µl of Conditioned medium+ 2.5 µl of TransDux+ 100 µl Max Enhancer) using TransDux™ MAX Lentivirus Transduction Reagent (#LV860A-1, System Biosciences). After 72 h, stable cells were selected using Puromycin (1µg/ml).

**Glycoprotein Isolation**

Glycoproteins were isolated using a glycoprotein isolation kit (89804; Thermo Fisher Scientific) as per the manufacturer’s protocol. Briefly, endothelial cell lysates were prepared using RIPA buffer (BP-115, Boston Bioproducts) containing protease inhibitor cocktail (#5872, Cell Signaling Technologies, 1x) without EDTA or other metal chelators. ConA Lectin Resin was washed in 1x Binding/Wash buffer (twice) by centrifugation of a column containing 50% resin slurry (200 µl per sample) for 1 min at 1000x g. Endothelial cell lysates (1000 µg) were added to the resin and incubated for 10 min at room temperature with end-to-end mixing (capped columns) using a rotator. Columns were centrifuged to separate resin bound to glycosylated protein and unbound lysates (flow through). Resins were washed in 400 µl of 1x Binding/Washing buffer twice (5 min incubation each) in the column. Bound glycosylated proteins were eluted from resins by adding 200 µl Elution buffer for 5 min at room temperature (with end-to-end mixing) followed by centrifugation at 1000x g for 1 min. Glycosylated, un-glycosylated (flow through) and total proteins were quantified and 50 µg of proteins were loaded on SDS PAGE gel for assessing the level of FcRn and Sparc glycosylation using their antibodies.

**Co-immunoprecipitation (Co-IP)** Protein–protein interactions between Galectin-3 and FcRn were studied using Co-Immunoprecipitation Kit (ab206996, Abcam). Mouse endothelial cells (bEnd.3) were lysed in cold lysis buffer (non-denaturing) and 800 μg of protein of different lysates were incubated with 5 μg of Galectin-3 and FcRn primary antibody (α-Galectin-3: ab76245 and α-FcRn: LS-C408079) overnight on a rotary mixer at 4°C. Protein A/G Sepharose beads was washed with 1× wash buffer (twice) and resuspended as 50% slurry. Following antibody binding, Protein A/G Sepharose beads (30 μl per sample) was added to each tube and incubated for 1 hour at 4°C. The beads were washed thrice with wash buffer and were finally collected by centrifugation (2,000 × g for 2 minutes at 4°C). Bound proteins/ complexes were eluted by boiling (5 minutes) the beads in denaturing 2× SDS-PAGE loading buffer. Samples were resolved on Any KD Mini-PROTEAN TGX (Bio-Rad, catalog no. 456-9033) SDS-PAGE gel and processed for Western blot analysis. As control, 5% of total lysate used in the assay was loaded on a separate gel.

**Colocalization using confocal microscopy.** Mouse endothelial cells (bEnd.3) were plated (2x10^4^ cells/well) on chamber slides and allowed to grow for 24 hours. Cells were washed with PBS and fixed in 4% PFA for 10 min. Cells were then washed twice with PBS and blocked in 5% Goat serum (Vector Laboratories #S-1000) containing 0.2% Triton X-100 in PBS at RT for 1 hr. Cells were incubated with primary antibody (α-FcRn antibody LS-C408079, LSBio, dilution 1:50; α-Galectin-3 antibody, MA1-940, Thermo Fisher Scientific, dilution 1:100) overnight at 4°C. Cells were washed thrice in PBS for 2 min each and were incubated with secondary antibody in 5% goat serum (Alexa Fluor™ 594 nm goat-anti-mouse and Alexa Fluor™ 488 nm goat-anti-rabbit, Thermo Fisher Scientific, dilution 1:2,000) at RT for 60 min in the dark along with DAPI (1:500). Cells were washed in PBS thrice and chamber slide was mounted with a coverslip using mounting media and visualized in confocal microscope.

**Immufluorescence of frozen human brain metastasis specimens.**

Nine flash frozen human craniotomy specimens were collected at the Massachusetts General Hospital Cancer Center, Harvard Medical School (IRB 10-454), the Military Institute of Medicine and the Copernicus Hospital Gdańsk in Poland (Bioethics committee approval number: 52/WIM/2015), and the Hospital Clairval, and stored in the AP-HM tumor bank (AC-2013-1786), in France. Informed consents were obtained from all the subjects for the samples. All the samples were anonymized and approved by the Office of Human Subjects Research Protections (OHSRP) at the National Institutes of Health (OHSRP #13093). IF staining was performed as described previsouly^19.^ Briefly, tissues were embedded in OCT. Slides were fixed with methanol for 5 min at -20°C, washed with PBS then incubated in blocking buffer (PBS with 5% goat serum (DAKO) for 20 min at room temperature. Primary antibodies to Galectin3 (LSBio cat# LS-B6768, 1/50), neonatal Fc receptor (FcRn) (Invitrogen cat# A5-42871, 1/50), LRP1 (Abcam, cat#ab92544, 1/100), SPARC (LSBio cat#LS‑B11553., 1/50) Collagen IV (Millipore cat# ab756P, 1/100) and/or Human Cytokeratin (DAKO clone MNF116, 1/100) were incubated overnight at 4°C. After 3 washes, the secondary antibodies (1:500, Alexa fluor® antibodies) and DAPI were incubated for 1 hr at room temperature. The slides were mounted using fluorescence mounting medium (Dako). Pictures were acquired with Zeiss Axioskop and ZEN software^19^.

**Immunohistochemistry (IHC) of human brain metastasis specimens.** Formalin-fixed, paraffin embedded blocks of human craniotomy specimens originating from breast, lung and ovarian cancers were collected by the Military Institute of Medicine and Medical University of Gdańsk in Poland (Bioethics committee approval number: 52/WIM/2015). Informed consents were waived as the patients were deceased.

All procedures were performed according to the manufacturer’s instructions (LifeSpan, Inc.). Tissue sections were deparaffinized in xylene and rehydrated through graded alcohol concentrations (100%, 96%, 80%, and 70%). For antigen retrieval, slides were pre-treated with a low pH target retrieval solution (Dako K8005). Endogenous biotin was blocked with an appropriate kit. Sections were incubated for 1 hour with an antibody against the human Galectin-3 (Monoclonal Rat anti Mouse LGALS3 / Galectin 3 Antibody (clone M3/38, LSB6768, 1:100 dilution). The antigen–antibody complex was visualized using the ImmPRESS Goat Anti-Rat IgG (Mouse adsorbed) Polymer Kit MP-7444 (VECTOR). Colon tissue served as a positive control. The immunoreactivity was scored semi-quantitatively on a 0-3+ intensity basis in the endothelial, cancer, and brain parenchymal compartments. Data are shown as percentage of specimens with any staining, or by staining intensity.

**Graphic representation and statistical analysis**. The data were graphed and analyzed using the software GraphPad Prism 7.01 (GraphPad Software, Inc.). Specific tests are mentioned in each figure legend. Non-parametric tests were performed when the data were not normally distributed. For each replicate, multiple measurements were recorded at each time point. The exact number (n) of replicates used in each experiment are reported in the respective figure legends. Statistical differences were calculated using One-way analysis of variance (ANOVA) using Tukey's multiple comparisons test. Each P value is adjusted to account for multiple comparison and 95% confidence interval (0.05) was used to determine the significance. For animal studies, animals were randomly assigned to the different experimental groups and experimenters were blind regarding group assignments.
