## Supplementary Fig. S1-S8 for "Comparison of Three Transcytotic Pathways for Distribution to Brain Metastases of Breast Cancer"

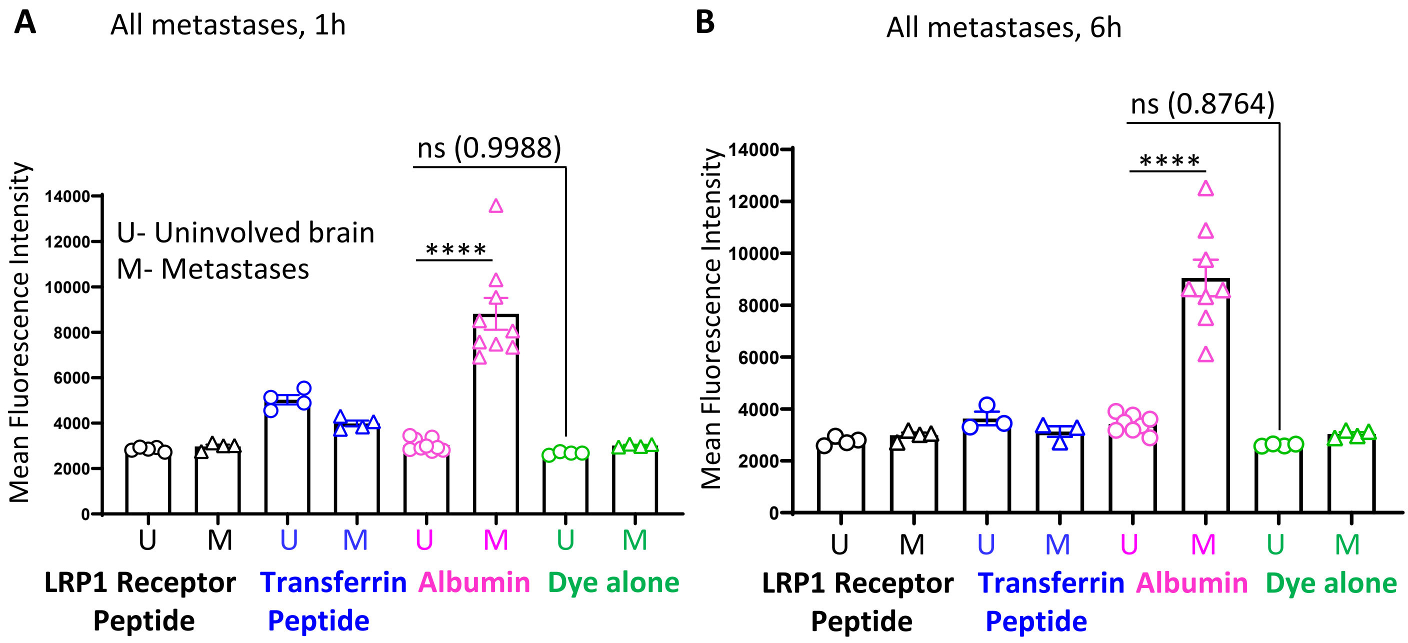


**Supplementary Fig S1. Comparison of three transcytosis pathways for uptake in the HER2+ JIMT1-BR experimental brain metastasis model of breast cancer.** Experimental design same as Figure 1 using JIMT1-BR HER2+ brain-tropic breast tumor cells (JIMT1-BR). Whole brain sections were scanned for eGFP and Far-red intensity. Mean fluorescence intensity of each probe in uninvolved (metastasis free, U) brain and brain metastases (M), in all metastases at two circulation timepoints 1h (A) and 6h (B) are plotted. Statistical differences were calculated using One-way analysis of variance (ANOVA) by comparing uninvolved brain (U) to metastases (M) across all the groups. ****, $P < 0.0001$

### Supplementary Fig. S2

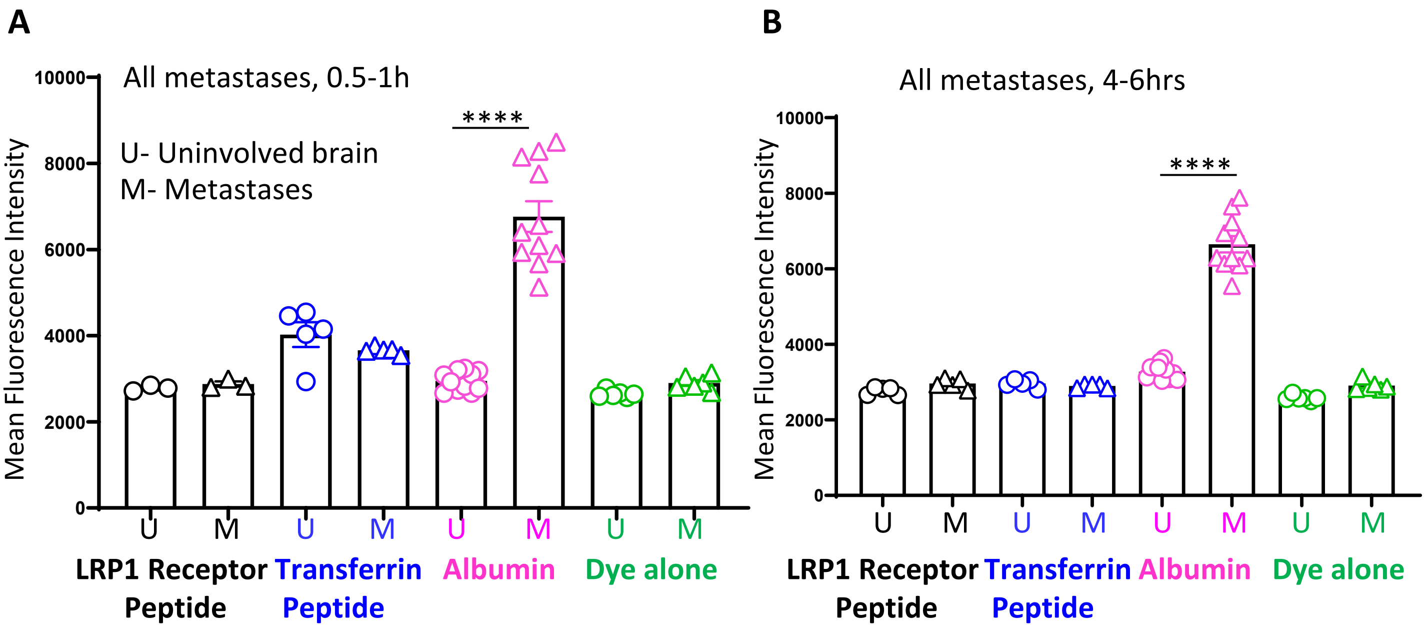


**Supplementary Fig S2. Comparison of three transcytosis pathways for uptake in the triple-negative 231-BR experimental brain metastasis model of breast cancer.** Experimental design same as Figure 1 using triple-negative 231-BR brain-tropic breast tumor cells (231-BR). Whole brain sections were scanned for eGFP and Far-red intensity. Mean fluorescence intensity of each probe in uninvolved (metastasis free, U) brain and brain metastases (M), in all metastases at two circulation timepoints 0.5-1h (A) and 4-6hrs (B) are plotted. Statistical differences were calculated using One-way analysis of variance (ANOVA) by comparing uninvolved brain (U) to metastases (M) across all the groups. ****, $P < 0.0001$

Supplementary Fig. S3

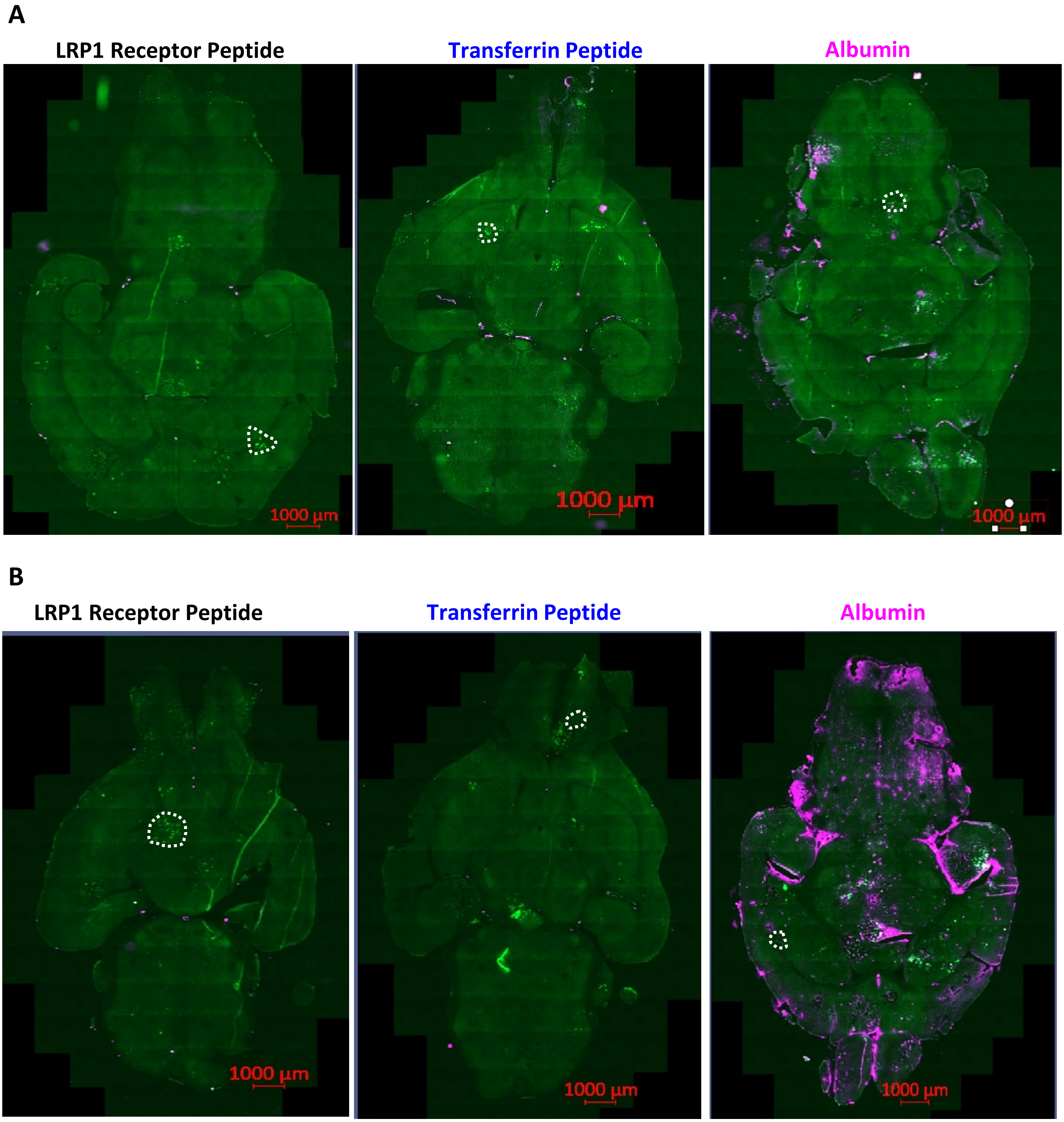


**Supplementary Fig S3. Photomicrographs of three transcytosis pathways for uptake in the triple-negative 231-BR experimental brain metastasis model of breast cancer.** Experimental design same as Figure 3 using triple-negative 231-BR brain-tropic breast tumor cells (231-BR). Representative merged (eGFP and Far-red-Albumin) photomicrographs of whole brain sections are shown for each transcytosis ligand at two circulation timepoints 0.5-1h (A) and 4-6hrs (B).

**Supplementary Fig. S4**

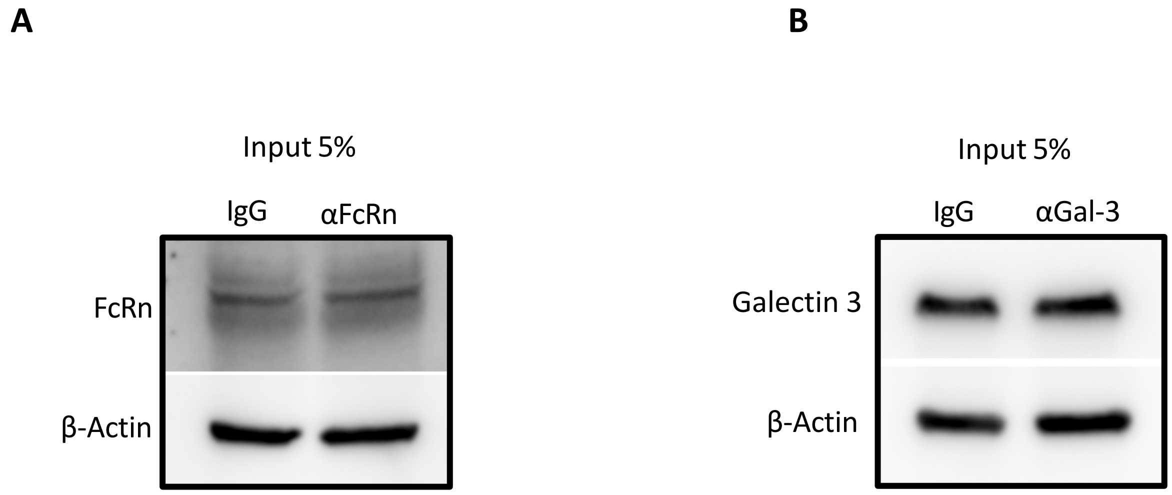


**Supplementary Fig. S4. Input controls for co-immunoprecipitation.** (A-B) Protein–protein interactions between Gal-3 and FcRn were studied using Co-Immunoprecipitation assay. Mouse endothelial cell (bEnd.3) lysates were incubated with FcRn or Galectin-3 primary antibody overnight on a rotary mixer at 4°C. Antibody bound proteins/complexes were isolated using Protein A/G Sepharose beads. Samples were resolved on Any KD Mini-PROTEAN TGX. As control, 5% of total lysate used in the assay was loaded on the above gel as input and processed for Western blot analysis

**Supplementary Fig. S5**

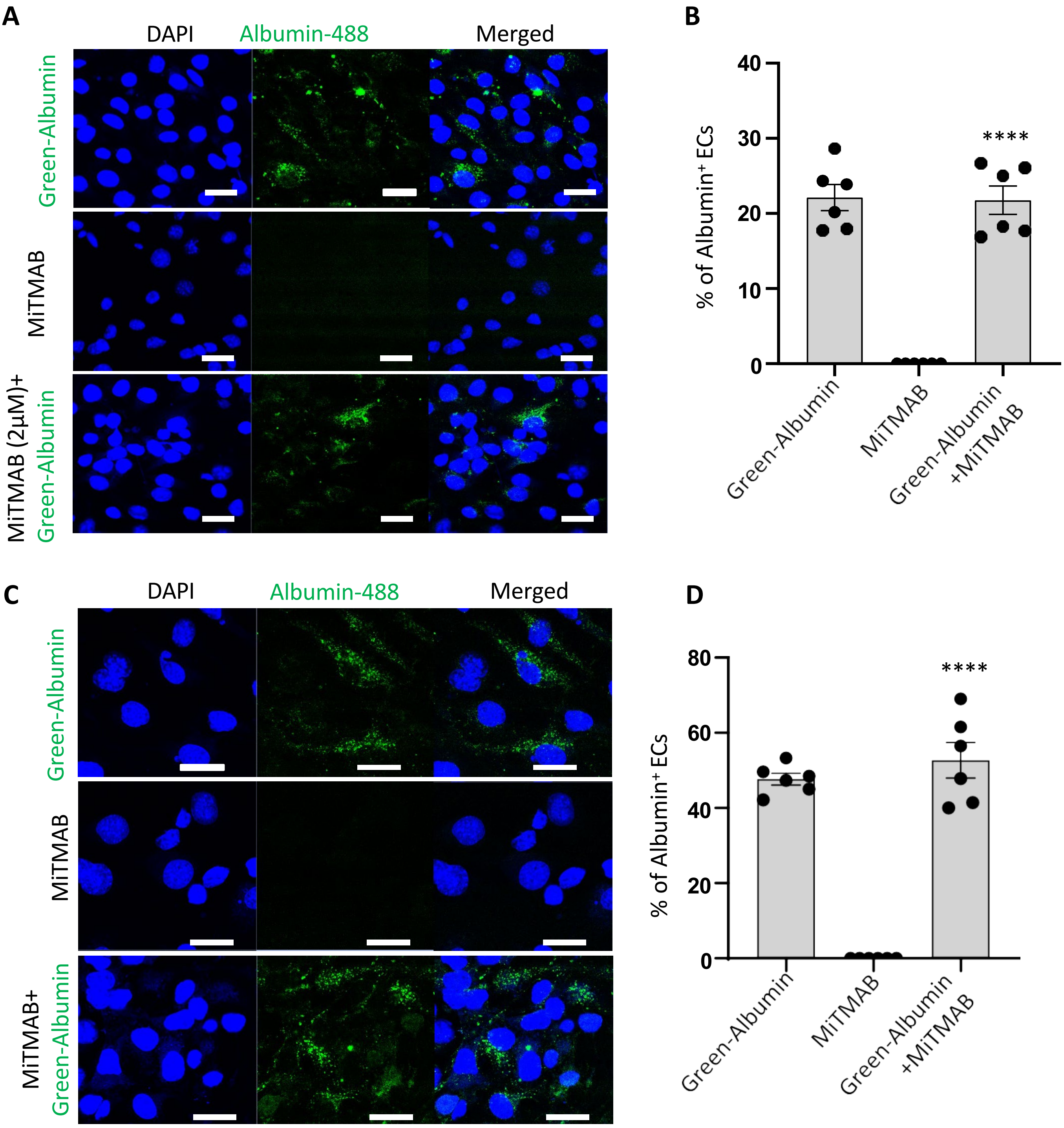


**Supplementary Fig. S5 Green-Albumin uptake by endothelial cells is dynamin independent.** (A-D) Endothelial cells were treated with 2 µM of MiTMAB for 30 min, an inhibitor of Dynamin dependent endocytosis, before addition of 50 µg/ml Albumin-488, which was incubated for 30 min. Endocytosis was determined using an *in vitro* BTB assay with 231BR (A, B) and JIMT1-BR (C, D) cells using confocal microscopy and quantitated (n=6). Scale bar 10 µm. All experiments were performed at least six times and statistical differences were calculated using One-way ANOVA.

### Supplementary Fig. S6

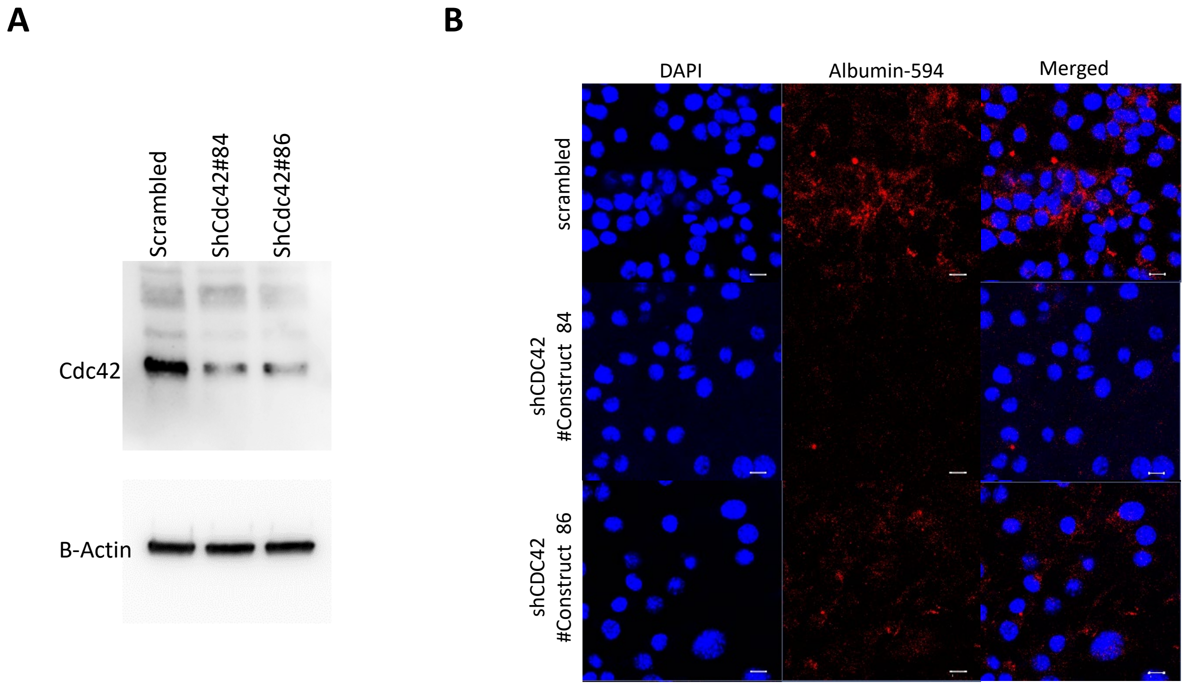


**Supplementary Fig. S6. Endothelial knockdown of Cdc42 disrupts albumin endocytosis.** (A) Cdc42 was knocked down in mouse brain endothelial cells using two independent shRNAs and compared to a scrambled shRNA by western blot. (B) Representative confocal images of endothelial cultures used for albumin endocytosis in *in vitro* BTB assay are shown. Scale bar 20 µm.

### Supplementary Fig. S7

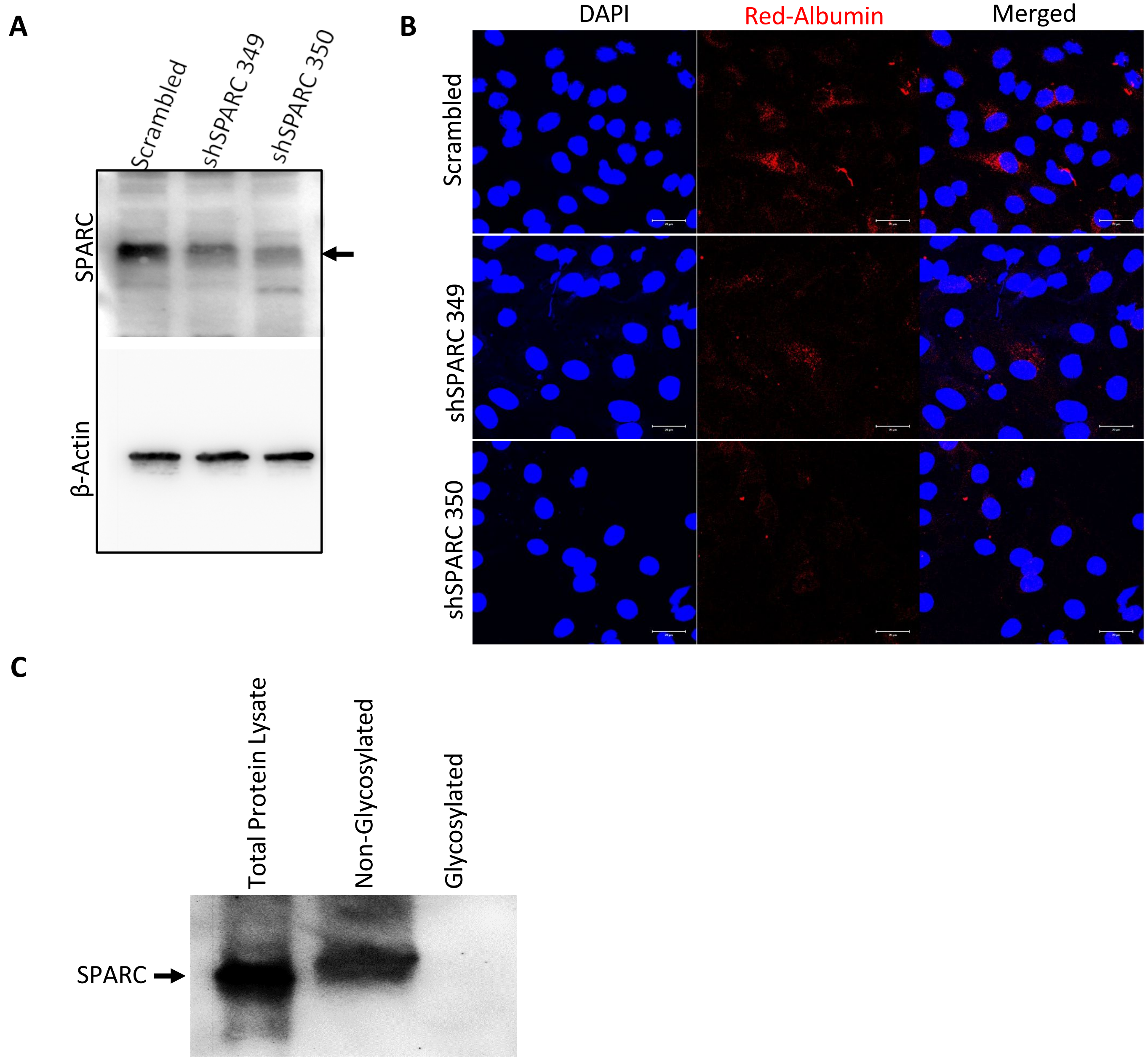


**Supplementary Fig. S7. SPARC knockdown showed heterogeneous albumin uptake in clones and is non-glycosylated. (A)** SPARC was knocked down in mouse brain endothelial cells using two independent shRNAs compared to a scrambled shRNA as shown by Western blot (band is indicated by an arrow). (B) Endothelial cultures were used for albumin endocytosis in the BTB assay on JIMT1-BR cells. A representative confocal images with heterogenous staining in both clones is shown; scale bar 20 µm. (C) Brain endothelial cell lysate was separated into glycosylated and non-glycosylated fractions using Concanavilin A based glycoprotein isolation kit and probed for SPARC (band shown by an arrow). No glycosylation band was seen in SPARC.

### Supplementary Fig. S8

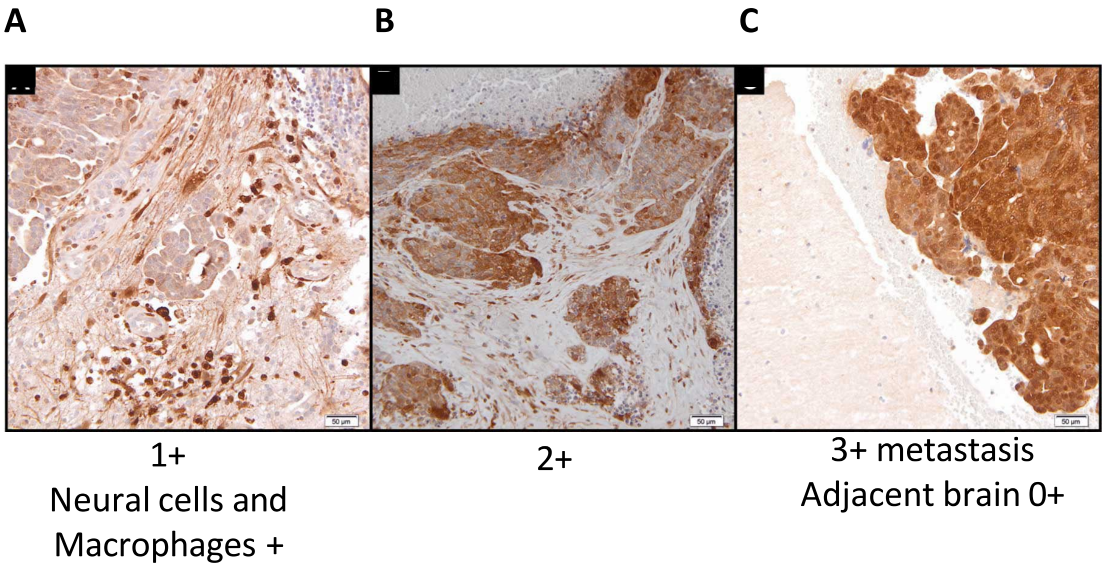


**Supplementary Fig. S8. Representative immunohistochemical intensity staining**. Sections from formalin-fixed, paraffin embedded human craniotomy specimens were stained for Gal-3 and analyzed by a pathologist. The intensity scale was used to quantify staining: 0 to 3+. Examples of staining are shown **A**-**C**. Magnification bars 50 µm.
